# Extensive hidden genetic variation shapes the structure of functional elements in *Drosophila*

**DOI:** 10.1101/114967

**Authors:** Mahul Chakraborty, Roy Zhao, Xinwen Zhang, Shannon Kalsow, J.J. Emerson

## Abstract

Mutations that add, subtract, rearrange, or otherwise refashion genome structure often affect phenotypes, though the fragmented nature of most contemporary assemblies obscure them. To discover such mutations, we assembled the first reference quality genome of *Drosophila melanogaster* since its initial sequencing. By comparing this genome to the existing *D. melanogaster* assembly, we create a structural variant map of unprecedented resolution, revealing extensive genetic variation that has remained hidden until now. Many of these variants constitute strong candidates underlying phenotypic variation, including tandem duplications and a transposable element insertion that dramatically amplifies the expression of detoxification genes associated with nicotine resistance. The abundance of important genetic variation that still evades discovery highlights how crucial high quality references are to deciphering phenotypes.

## Introduction

Mutations underlying phenotypic variation remain elusive in trait mapping studies (Rockman 2012) despite the exponential accumulation of genomic data, suggesting that many causal variants are invisible to current genotyping approaches (Eichler, et al. 2010;Manolio, et al. 2009;McCarthy, et al. 2008; Wray, et al. 2013). Moreover, despite contributing substantially to genome sequence variation, mutations affecting genome structure, like duplications, deletions, and transpositions (Alkan, et al. 2011a; Emerson, et al. 2008), are systematically underrepresented by standard methods (Alkan, et al. 2011a), even as a consensus emerges that such structural variants (SVs) are important factors in the genetics of complex traits (Eichler, et al. 2010). Addressing this problem requires compiling an accurate and complete catalog of genome features relevant to phenotypic variation, a goal most readily achieved by comparing nearly complete, high-quality genomes (Alkan, et al. 2011a). This standard was first achieved for metazoans in a scalable way with the completion of the *Drosophila melanogaster* genome (Myers, et al. 2000). The sequencing of *D. melanogaster* by whole genome shotgun sequencing (WGS) catalyzed an explosion of genome projects that aimed to catalog genes and identify mutations responsible for phenotypes. Subsequent development of high-throughput short-read sequencing led to an even steeper drop in cost and a commensurate increase in the pace of sequencing (2010). However, adoption of these methods led to a focus on single nucleotide changes and small insertion deletions (Frazer, et al. 2009; Wray, et al. 2013) and, paradoxically, a deterioration of the contiguity and completeness in new genome assemblies, due primarily to limitations in read length and fragment size (Alkan, et al. 2011b).

## Results and discussion

Here we present a reference quality assembly of a second *D. melanogaster* genome and introduce a comprehensive map of SVs that reveals a vast amount of hidden variation. Collectively, newly discovered SVs both exceed the total variation due to SNPs and small indels and include strong candidates for explaining phenotypic variation in mapped complex traits. We discovered these variants by comparing the existing genome of the *Drosophila melanogaster* strain ISO1 to a new high-quality reference-grade assembly of a cosmopolitan *D. melanogaster* strain from Zimbabwe called A4. The A4 strain is a part of the Drosophila Synthetic Population Resource (DSPR) (King, et al. 2012), a widely-used trait mapping resource that represents a model for discovery of phenotypically relevant variants. We assembled the new A4 genome using high coverage (147X) long reads using Single Molecule Real-Time sequencing on DNA extracted from females (Fig. S1). The A4 assembly is more contiguous than release 6 of the ISO1 strain — which is arguably the best metazoan WGS assembly — with 50% of the genome contained in contiguous sequences (contigs) 22.3 Mbp in length or longer (i.e. A4’s contig N50 is 22.3 Mbp; cf. ISO1’s N50 is 21.5 Mbp (Hoskins, et al. 2015); Table S1, Fig. S2-3). Compared to ISO1, the A4 assembly recovers more genome in far fewer sequences (144 Mbp in 161 scaffolds vs. 140 Mbp in 1,857 non-Y scaffolds) and exhibits an essentially identical level of completeness as measured by universal single-copy orthologs (Materials and Methods, Table S1) (Simao, et al. 2015). On a large scale, both genomes are co-linear across all major chromosome arms, making large-scale misassembly unlikely (Fig. 1a). Comparison of an optical map of the A4 genome and the A4 assembly confirms this inference by showing little evidence of misassembly introduced at either the assembly or scaffolding stage (Fig. S4-S5).

**Figure 1. a).**
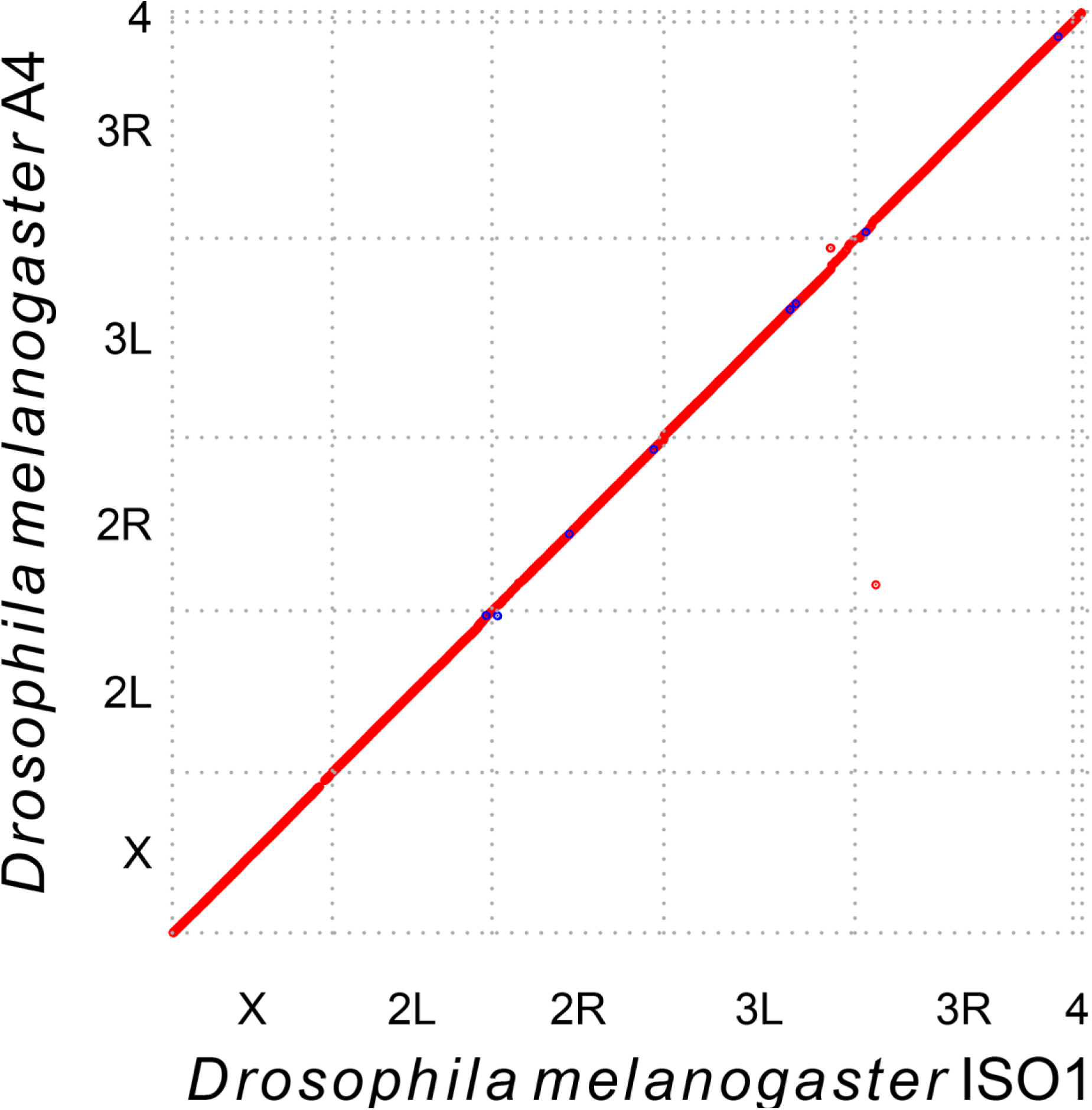
A dot plot between the reference (ISO1) chromosome arm scaffolds and the A4 scaffolds. The A4 assembly is as contiguous as the ISO1 assembly (scaffold N50 = = 25.4Mb vs 25.2Mb; Table S1). The repeats and transposable elements have been masked to highlight the correspondence of the two genomes.

Putative SVs were identified by classifying regions of disagreement in a genome-wide pairwise alignment between A4 and ISO1 assemblies as insertion-deletions (indels), copy number variants (CNVs), or inversions. Within the euchromatic portion of the genome (Table S2), we discovered 1,890 large (>100bp) insertion-deletions (Table S3; Fig. S6) affecting more than 7 Mbp of euchromatin sequence content between the two genomes. In contrast, small indels (<100bp) and SNPs affected only 1.4 Mbp (indels: 722 kbp; SNPs: 687 kbp). Of the large indels, 79% (1,486/1,890) are transposable element (TE) insertions. Although discovering TE insertions is possible with paired end short reads, a previously published catalog of TE insertions in A4 based on 70X short-read coverage failed to find 37% of the TE insertions in A4 reported here (Cridland, et al. 2013) (Fig. 1b, Fig. S7, Table S4,). These insertions invisible to short-read approaches often occur when a TE is inserted near an existing TE (e.g. Fig. S8), presumably resulting in complex multiply mapping reads that are more difficult to interpret than simple insertions. One such complex insertion in A4 affects *Multidrug-Resistance Like Protein 1* (*MRP*), which is a candidate gene for resistance to chemotherapy drug carboplatin (King, et al. 2014) (2L:12,753,668-12,753,672; Fig. S8).

**Figure 1. b).**
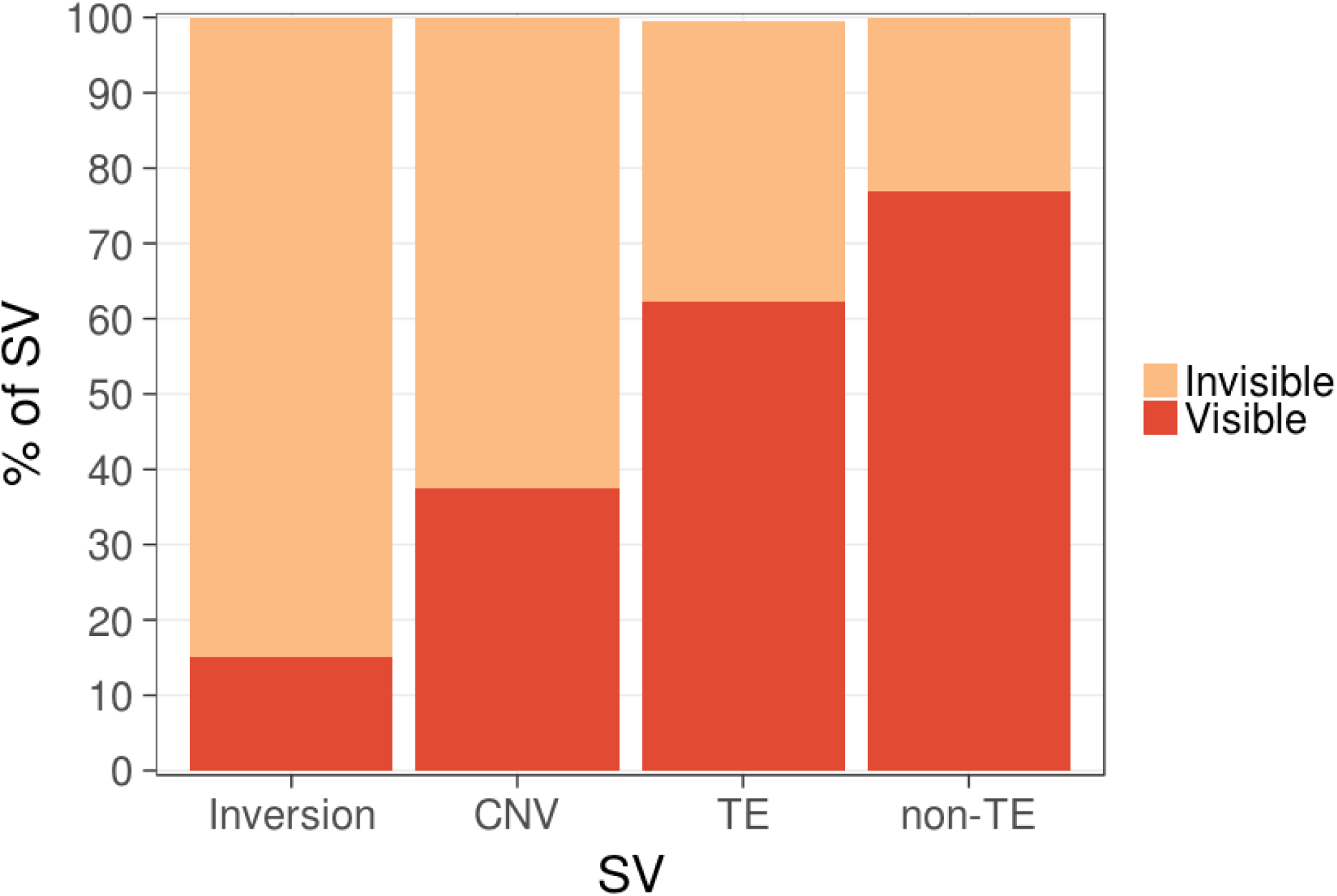
Proportion of large (>100bp) SVs in A4 chromosome 2L assembly that are not detected by SV genotyping based on paired end short reads. Illumina short-reads based TE indel genotypes were obtained from (Cridland, et al. 2013). For CNV and inversions, short-read based genotypes were obtained from the most reliable genotyping strategy (Materials and Methods).

A large proportion of TE insertions affect introns (395/718 in ISO1,435/768 in A4), often introducing dramatic increases in intron length (Fig. 1c; Fig. S9). This is perhaps not surprising, given that insertions into exons often disrupt genes (Table S5). Additionally, TEs inserted into exons can be spliced out, effectively becoming new introns. We see evidence of this in cDNA from ISO1 (Stapleton, et al. 2002) and RNAseq reads in A4 that span what are large (>1kb) TE insertions into exons in the other genome (Table S5; Fig. S10-12). This provides evidence for gain of novel polymorphic introns via TE insertions (Table S5) and represents the first genome-wide glimpse of TE-derived introns segregating in a population. We discovered putative polymorphic TE introns both in genes of unknown function (e.g. *CG33170* in ISO1 or *CG13900* in A4) as well as into a well-understood developmental gene *(Polycomb* in ISO1; Table S5). TE insertions within introns are associated with decreased transcription (Cridland, et al. 2015), which may result from a phenomenon that slows transcription in long introns known as intron delay (Swinburne and Silver 2008). TE insertions that modulate the expression of important genes can have a direct impact on phenotype. Since most TEs have allele frequencies much less than 1% in *D. melanogaster* populations (Petrov, et al. 2011), not only are the number of bases in a genome affected by hidden TEs greater than the number affected by all SNPs and small indels combined (Table S3), they will be poorly tagged by common variants, complicating GWAS approaches for mapping traits.

**Figure 1. c).**
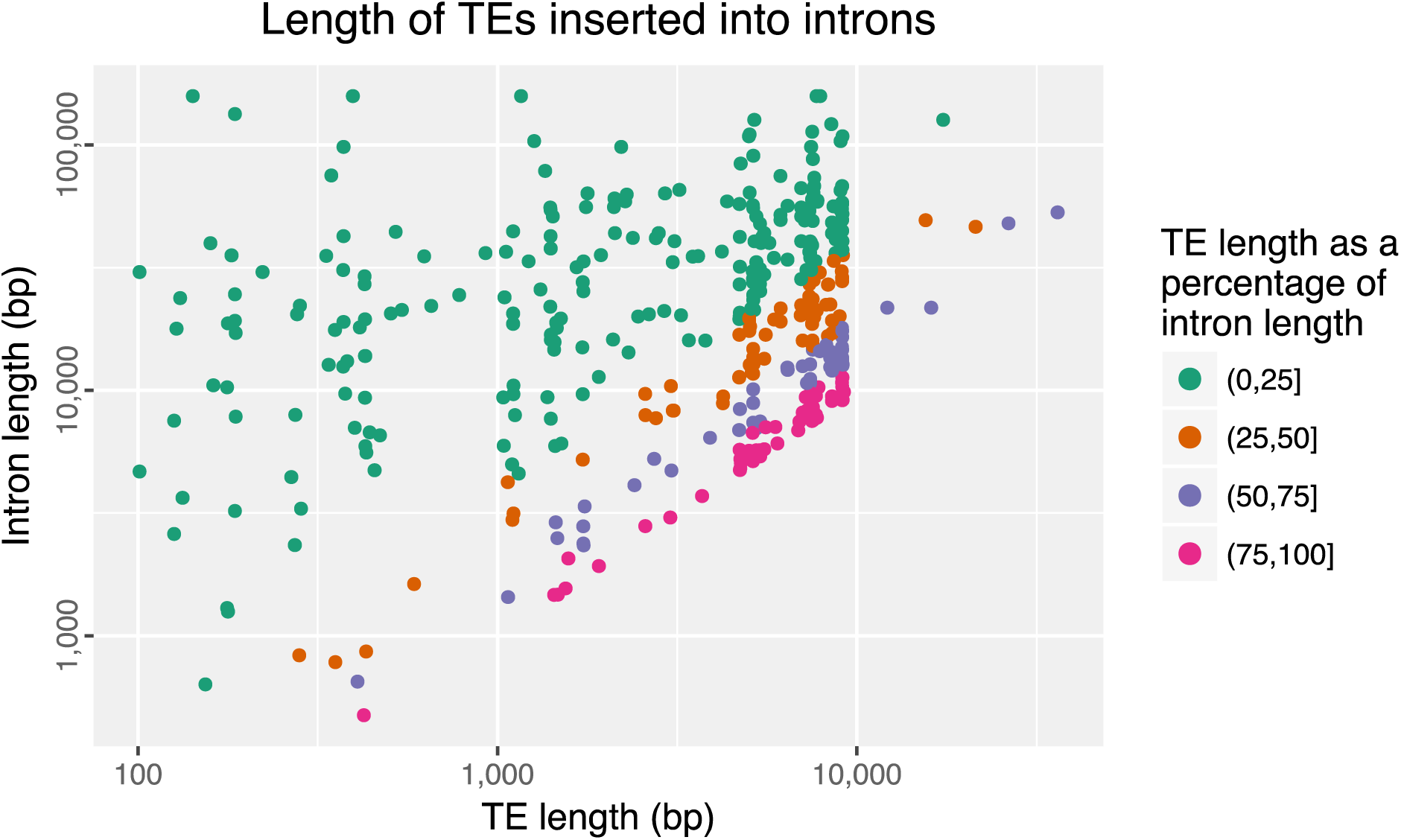
Relationship between polymorphic TEs length in ISO1 and the lengths of the introns they insert into. Most TEs are more than 1 kbp long (median 5.1 kbp). Many introns comprise mainly of TEs as evidenced by the insert sizes that are roughly equal to the intron lengths.

Non-TE indels in ISO1 and A4 represented 20% and 23% of the total number of such mutations, respectively, accounting for 170 kbp of sequence variation. On average, non-TE indels were much smaller than the TE indels (median 213 bp versus 4.7 kbp). Despite being small, 23% of these mutations could not be detected by paired end short reads (Fig. 1b). Nevertheless, non-TE indels often affect functional genes. For example, 18 genes have been partially deleted in A4 (Table S6). One of these genes, *Cyp6a17* is known to affect temperature preference (Kang, et al. 2011). Because exons 1 and 2 and intron 1 of the A4 *Cyp6a17* are deleted (Fig. S13), we predict temperature preference behavior of A4 differs from ISO1. Another deletion (129 bp) removed the second exon of a chitin binding protein gene called *Mur18B* (Fig. S14), and may contribute to protection from high temperature stress (MacMillan, et al. 2016). This deletion, which removes 41 amino acids from the Mur18B protein, likely renders the A4 allele of *Mur18B* a null mutant. However, despite this mutation being smaller than average short-read library fragment size, two different genome-wide deletion genotyping strategies based on short paired end reads failed to detect this mutation (Materials and Methods).

The A4/ISO1 comparison also uncovered 29 inversions, affecting a total of 60.6 kbp of sequence, ranging in size from 100 bp to 21 kbp (Table S3). Notably, only 4 of these inversions were detected by paired end short reads (see Materials and methods; Fig. 1b, Table S4). Despite their small numbers, inversions in our SV map often (21/29) affect regions harboring genes known to be functional, such as a 21 kbp segment that consists of a cluster of five gustatory receptor genes, including *Gr22a, Gr22b, Gr22c, Gr22d,* and *Gr22e* (Table S3). Interestingly, the A4 optical map revealed an additional large inversion that could not be resolved by the A4 assembly. This putative inversion occupies 300 kbp of the proximal end of the X chromosome scaffold (Fig. S4-5). Failure to resolve this inversion in A4 is not unexpected, because assemblies using a WGS approach tuned for euchromatin perform poorly in heterochromatic regions (Khost, et al. 2016).

We also detected 390 duplication CNVs (209 in A4 and 181 in ISO1) affecting ~600 kbp (Fig. 1d, Fig. S15, Table S3). We estimate that ~60% of these mutations are hidden from standard short-read detection methods (Fig. S16). Unlike indels, most CNVs affected exons (64%), with 34 duplicates encompassing full-length protein coding genes. Notably, among the 34 protein coding genes that are duplicated in A4, 13 were missed by short-read CNV genotyping methods (Materials and Methods). In total, only about ~40% of CNVs were discoverable with short-read methods exhibiting high specificity (Fig. 1b, Fig. S16), consistent with previous observations in mammalian genomes (Huddleston and Eichler 2016), preventing the discovery of many putative regulatory variants caused by CNVs. For example, a previous experiment compared the expression levels in larvae of the genes in A4 and another DSPR strain from Spain, called A3, to identify the gene regulatory changes underlying nicotine resistance (Marriage, et al. 2014). Interestingly, the comparison revealed 17 upregulated genes in A4 which are also duplicated in A4 (Table S7). Several of these genes have been previously identified as candidates for cold adaptation, variation in olfactory response, and toxin resistance, among others (fig 2a, 2b, Table S7-S8). Interestingly, eight of these CNVs were invisible to short read methods (Table S7), potentially misleading inferences about the mechanisms of regulatory variation.

**Figure 1. d).**
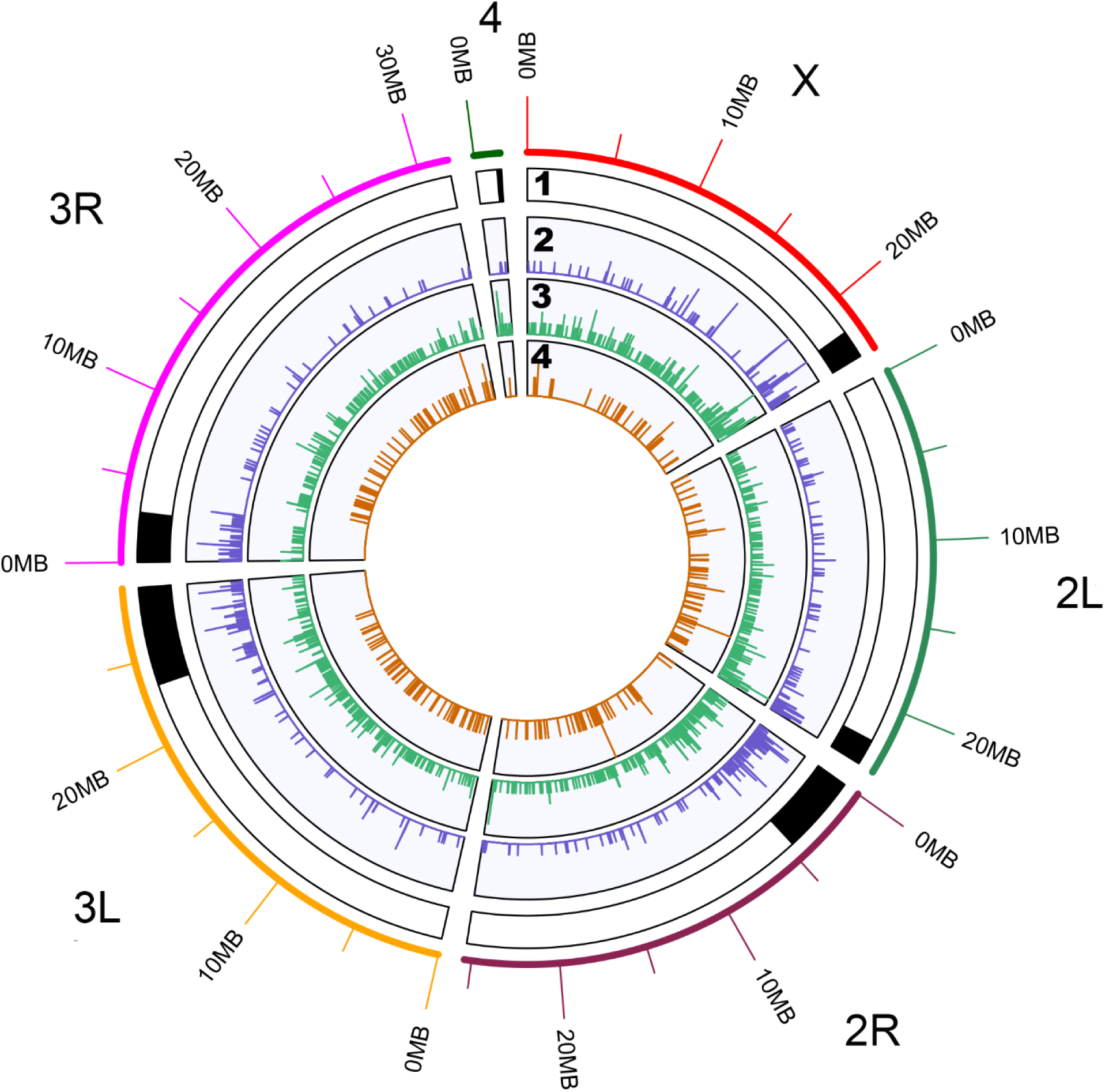
Distribution of SVs (>100bp) across the A4 chromosome arms. The segments shaded in black on track 1 are pericentric heterochromatin. Tracks 2-4 show SVs, including TEs, duplicate CNVs, and non-TE indels greater than 100 bp, respectively. CNVs and TEs are present in higher densities in heterochromatin, whereas non-TE indels are less numerous.

Among the eight upregulated hidden duplicates in A4, QTLs containing the genes *Cyp28d1* and *Ugt86Dh* have been associated with resistance to nicotine, one of a common family of plant defense toxins called nicotinoids (Glendinning 2002; Marriage, et al. 2014). One QTL (Q1), accounting for 8.5% of the variation in nicotine resistance, contains two cytochrome P450 enzyme genes, *Cyp28d1* and *Cyp28d2,* both of which are upregulated (Marriage, et al. 2014). The other major effect candidate region, Q4, explains 50% of the variation in nicotine resistance and contains the *Ugt86D* gene cluster which possesses several differentially regulated genes, including *Ugt86Dh* (Fig. 2c). In the nicotine breakdown pathway, the cytochrome P450 enzymes function upstream of the UDP-glucosyltransferase (Ugt) enzymes (Luque and O'Reilly 2002). Interestingly, neither the nicotine study nor SV genotyping approaches using A4 short-read data successfully identified the structural mutations we report here (Materials and Methods). Since the A4 larvae carry the high resistance alleles at both loci, we studied these newly discovered SVs at *Cyp28d1* and *Ugt86Dh* to determine whether they could explain the expression in A4.

In our *de novo* A4 assembly, the Q1 locus contains a 3,755 bp tandem duplication separated by a 1.5 kbp spacer region, creating two copies of the genes *Cyp28d1* (*Cyp28d1-p* and *Cyp28d1-d*) and *CG7742* (*CG7742-p* and *CG7742-d*) (Fig. 2a; Fig. S19-S20). While duplication can increase expression levels (Henrichsen, et al. 2009; Schmidt, et al. 2010), an extra gene copy alone is unlikely to cause the ~50-fold increase in expression level observed in absence of nicotine or the ~3-fold increase observed in the presence of nicotine (Marriage, et al. 2014). By calculating the paralog-specific expression levels of each *Cyp28d1* copy in A4 to that of the single copy *Cyp28d1* locus in A3 (Materials and Methods) we found that, in the absence of nicotine, *Cyp28d1-p* and *Cyp28d1-d* showed ~41-fold and ~6.3-fold higher expression in A4 relative to A3 (Fig. 2c) respectively, for a total of ~47-fold upregulation, similar to previous results (Marriage, et al. 2014). Inspection of the 1.5 kbp spacer sequence revealed it to be an insertion of a fragment spanning the 5’ end of a long terminal repeat (LTR) retrotransposon called *Accord* (Fig. 2a). The insertion of the *Accord* LTR upstream of another gene called *Cyp6g1* has been linked to upregulation of its Cytochrome P450 enzyme (Chung, et al. 2007), but the detailed mechanism of upregulation remains unknown. In the Q1 duplication, duplicates nearer *Accord* are more strongly affected than their more distal paralogs, with CG7742-d and *Cyp28d1-p* most strongly affected (Fig. 2a, 2c). Interestingly, the duplication plus *Accord* insertion at Q1 is also associated with ~10-15-fold upregulation of *Cyp28d2,* which was not duplicated. Such long range effect of the *Accord* insertion on the expression of these genes is consistent with local chromatin state changes observed in other LTR retrotransposon insertions (Rebollo, et al. 2012).

**Figure 2. a).**
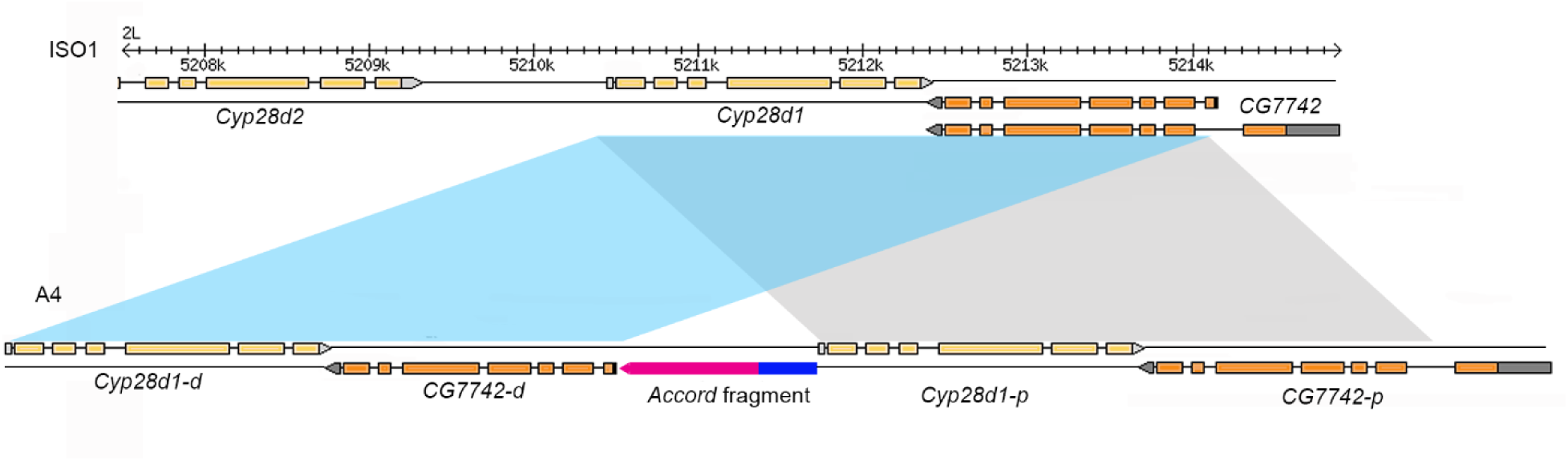
Duplication of *Cyp28d1* and *CG7742* in A4. The reference strain (ISO1) and A3 possess one copy of *Cyp28d1,* whereas A4 has two copies. A 1.5Kb *Accord* fragment (pink) containing an LTR (blue) is located between the proximal *Cyp28d1* and the distal *CG7742.* Grey rectangles denote UTRs and orange rectangles represent coding sequence.

The second nicotine resistance QTL, called Q4, contained several Ugt genes, including *Ugt86Dh.* Interestingly, higher expression of *Ugt86Dh* and *Ugt86Dd* in *D. melanogaster* has been implicated in increased resistance to DDT (Pedra, et al. 2004). Though a number of Ugt genes in Q4 show higher expression in the nicotine resistant A4 larvae than the nicotine sensitive A3 larvae (Marriage, et al. 2014) (Fig. 2b, Fig. S22-S23), candidate variants explaining these differences have yet to be identified. Interestingly, we find that *Ugt86Dh* is duplicated in A4 (Fig. 3a; Fig. S14), a mutation which remains undetected by paired-end short-reads (Table S4). However, unlike the *Cyp28d1* copies, each copy of *Ugt86Dh* is transcribed at similar levels, leading to a doubling of expression in A4 (Fig. 2c).

**Figure 2. b).**
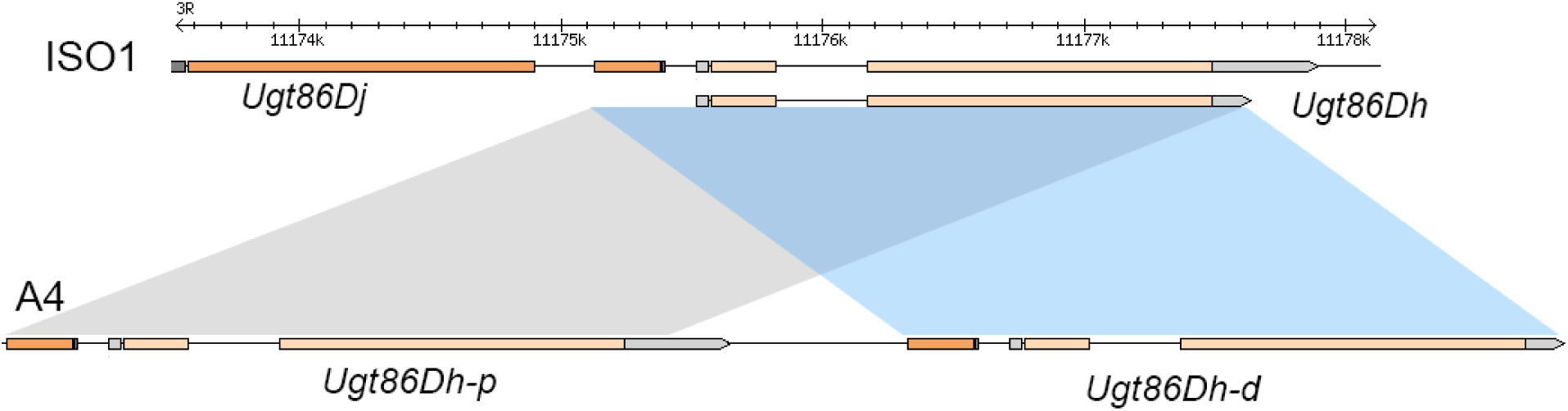
Tandem duplication of *Ugt86Dh* in A4. The duplication creates a new copy (*Ugt86Dh-d*) of an *Ugt86Dh* isoform consisting of a smaller 3’ UTR, and a copy of the first exon of the adjacent gene *Ugt86Dj.* A part of the *Ugt86Dh* first intron is also deleted in *Ugt86Dh-d*. Figure 2. a-b) The shaded parallelograms indicate the span of the duplicated segments, with gray representing the proximal copy, and blue representing distal copy with respect to the centromere.

**Figure 2. c).**
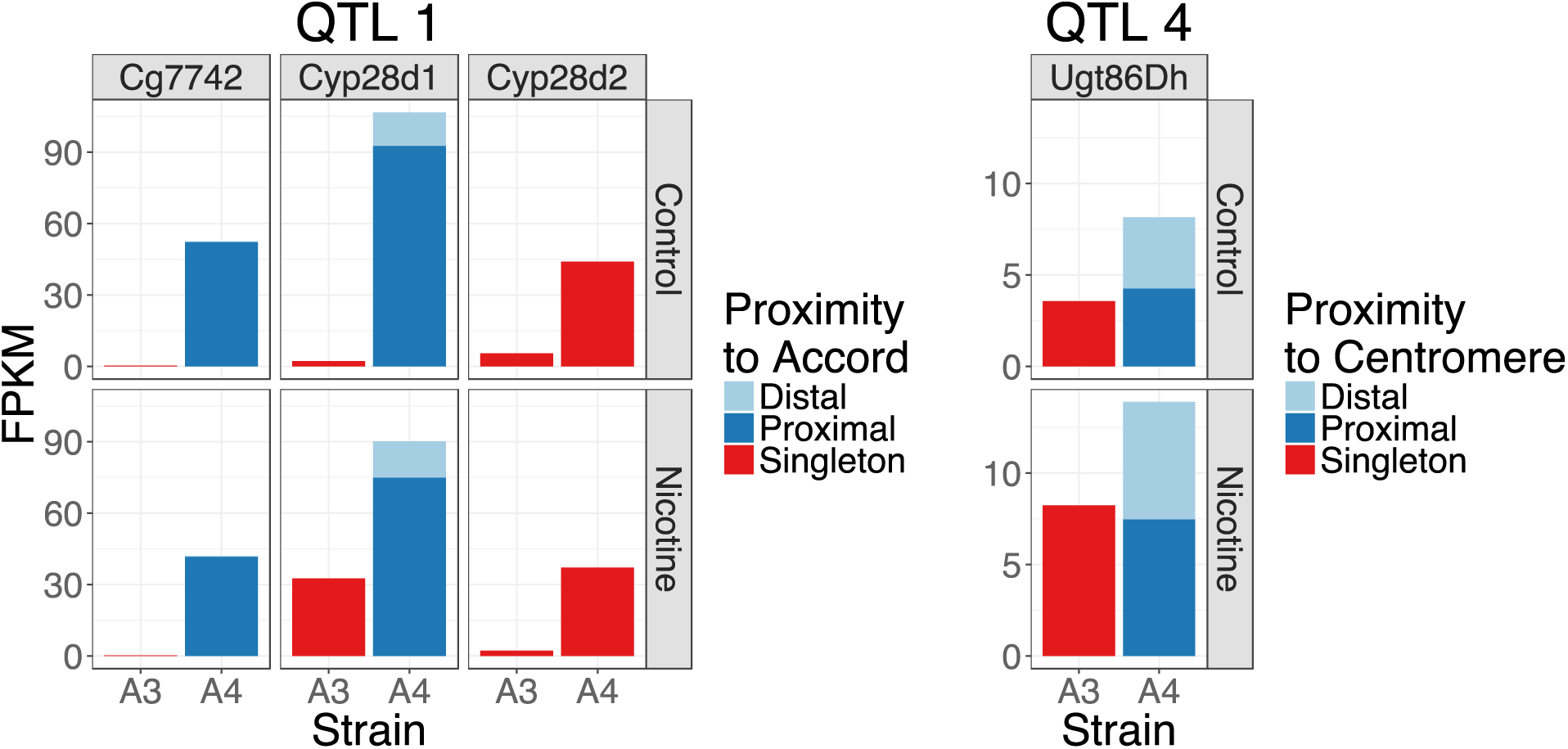
Paralog specific expression level of the Q1 (left) and Q4 (right) candidate genes in in A4 and A3 strains in presence and absence of nicotine in the food. Among the duplicated genes *CG7742* and *Cyp28d1,* the copies located nearer the *Accord* element are transcribed at higher levels than those located further away. While *Cyp28d1* upregulation in A4 is a combination gene duplication and TE insertion, *Cyp28d2* is likely explained by the *Accord* insertion. Unlike the duplicates at Q1, at Q4 both copies of *Ugt86Dh* are expressed at similar levels. When nicotine is present in the food, the expression level of the both gene copies nearly doubles.

**Figure 3. a).**
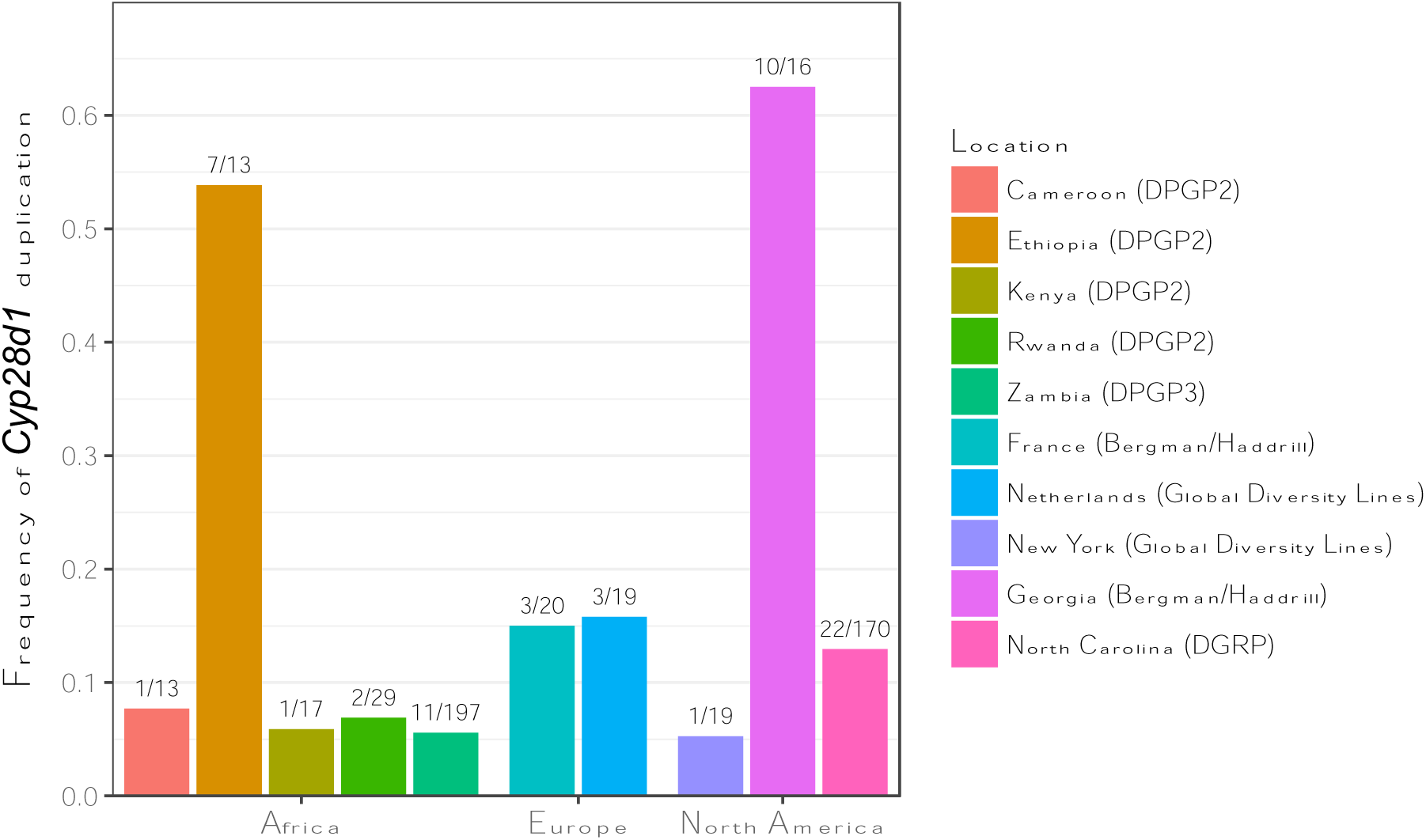
Combined frequency of four *Cyp28d* duplicate alleles in different population samples. The duplicates segregate at particularly high frequencies in Ethiopia and Georgia and at intermediate frequencies in North Carolina and Netherlands.

**Figure 3. b).**
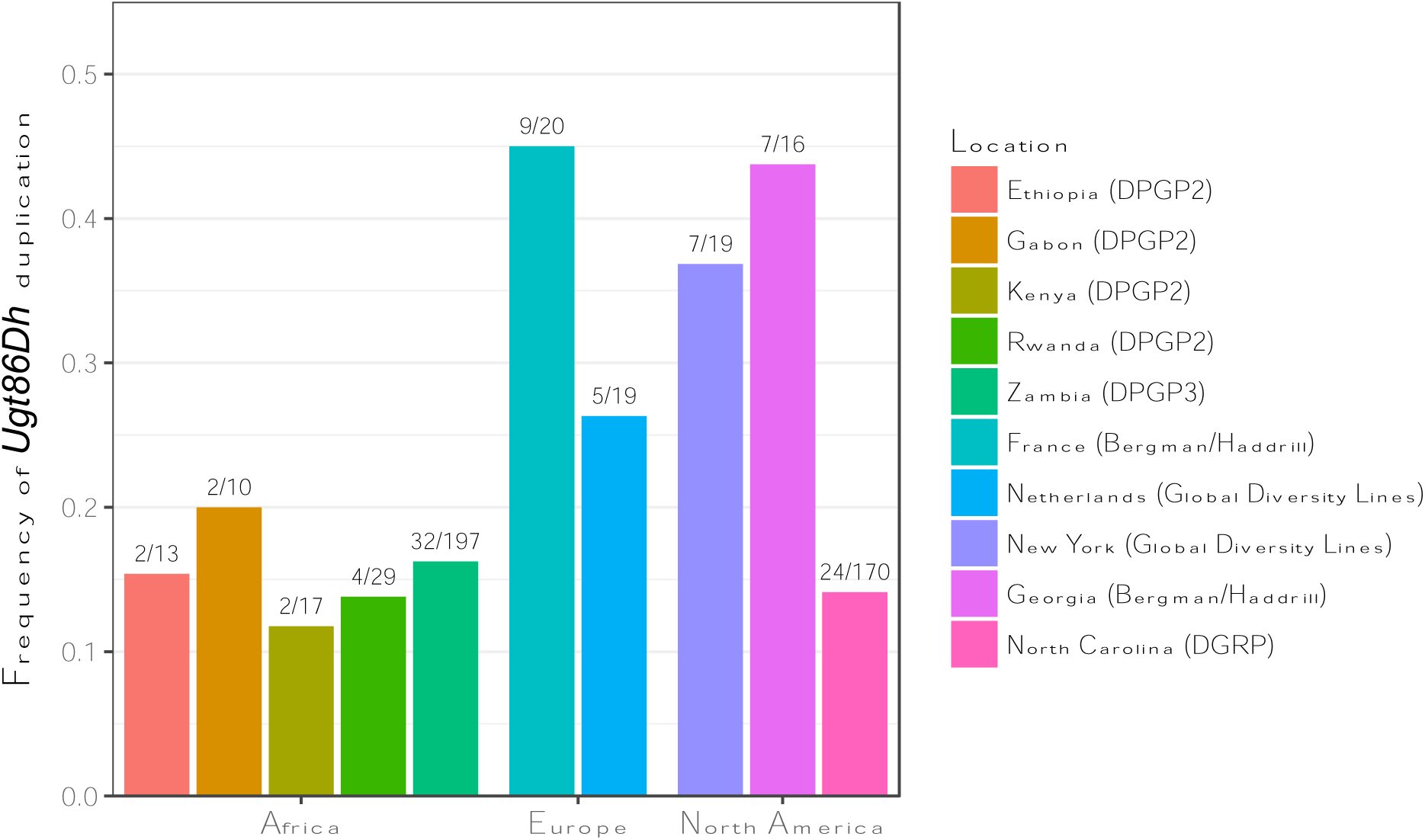
Frequency of the *Ugt86Dh* duplicate in different populations. *Ugt86Dh* duplicates are found in intermediate to high frequencies in all populations, with slightly higher frequencies in Europe and North America.

Like DDT, nicotine and its analogs have been widely used as pesticides. Hence, given the abundance of nicotinoids in the environment, we predict that mutations conferring resistance to nicotine would also be common. Consistent with this prediction we have found that duplicates encompassing *Cyp28d1* and *Cyp28d2* segregate at intermediate or high frequencies in multiple populations (Fig. 4a), all in a region spanning less than 25 kbp. Interestingly, these mutations include at least four alleles, which is remarkable given that the rate of SV heterozygosity between A4 and ISO1 in an average 25 kbp window is only 0.08. Additionally, the *Ugt86Dh* duplicate also segregates at high or intermediate frequency in nearly all *D. melanogaster* populations that we examined (Emerson, et al. 2008) (Fig. 4b). However, unlike for *Cyp28d1* and *Cyp28d2,* the Ugt86Dh mutations comprise only a single allele. Interestingly, patterns of SNP variation suggest recent bouts of natural selection in both regions exhibiting structural variation and regulatory variation associated with nicotine resistance (Figure S18 and S19).

So far, we have focused on novel variants discovered in A4, particularly those that were previously inaccessible to existing genotyping approaches. However, there are virtually identical numbers of variants in ISO1 and A4. There is no biologically meaningful sense in which ISO1 is a more appropriate reference than any other strain. Projects like ENCODE and modENCODE (mod, et al. 2010) have expended substantial effort annotating reference genomes of one genotype with functional genomic data obtained from different genotypes or cell lines. Without high quality reference genomes associated with the experimental genotypes, rare mutations segregating in the reference will result in errors in inference. In particular, approaches like RNAi or CRISPR require precise sequence information about their targets that can be easily misled by hidden SV. For example, a study about the origin of new genes in *Drosophila* made the remarkable claim that new genes rapidly become essential (Chen, et al. 2010). This study reported one putative essential gene, a p24 transporter called *p24-2,* so young that it is present only in *D. melanogaster.* Experiments aimed at knocking out this gene using RNAi constructs suggested that although new, *p24-2* is essential. However, though present in ISO1, *p24-2* is absent in the A4 assembly (Fig. S23), and is likely also absent in the strains used to carry out the functional work. That this new gene is absent in a healthy strain like A4 refutes the essential status of this gene in *D. melanogaster.*

The ubiquity of hidden variation in genome structure is merely a first glimpse beneath the tip of an iceberg of genetic variation governing phenotypes. In concert with careful phenotypic measurements, a new wave of high quality genomes will reveal heritable phenotypic variation invisible to short-read approaches, like those caused by structural mutations including transposable elements, duplications, and repeats, among others. While previous estimates on the relative contributions of SVs and SNPs toward regulatory variation suggested that the former is modest (Stranger, et al. 2007), our results show that popular genotyping approaches miss a significant number of SVs (Fig. 1b, Fig. S7,S16, Table S4), including those which impact gene expression and organismal phenotype (Table S7-S8). Consequently, previous estimates of the contribution of SVs towards regulatory and phenotypic variation may be misleading (Gamazon, et al. 2011). The large fraction of hidden variation we report here is based on only the euchromatin portion of *D. melanogaster,* a species likely harboring fewer complex structural features than other higher eukaryote model systems or other animal and plant species important in food production. Our results suggest that the medical and agricultural impact of hidden variation is likely much greater than previously appreciated in systems such as humans, and crop species like wheat and maize.

## Materials and Methods

### DNA sequencing

A4 DNA was extracted from females following the protocols described in (Chakraborty, et al. 2016) and the raw genomic DNA was sheared using 10 plunges of 21 gauge needle followed by 10 pumps of the 24 gauge needle. SMRTbell template library was prepared following the manufacturer’s guidelines and sequenced using P6-C4 chemistry in Pacific Biosciences RSII platform. We sequenced 30 SMRTcells corresponding to 19.1 Gb (50% of the sequences are contained within 18Kbp or longer reads) of nucleotide sequences. All sequencing was performed at University of California Irvine Genomics High Throughput Facility.

### Genome assembly

To assemble the genome, the pipeline described in (Chakraborty, et al. 2016) was followed. For all calculations of sequence coverage, a genome size of 130Mbp was used (G =130×10^6^bp). We generated a hybrid assembly (NG50 =4.23Mbp; assembly size=129Mbp) with *DBG2OLC* (Ye, et al. 2016) using the longest 30X PacBio reads and 74.6X paired end Illumina short reads from King et al. (King, et al. 2012). The PacBio only assembly (NG50 =13.9Mbp; assembly size = 147Mbp) was generated using PBcR-MHAP (Berlin, et al. 2015) pipeline as implemented in *wgs 8.3rc1.* Next, *quickmerge* (v.0.1, parameters hco =5, c= 1.5, l = 2Mb)(Chakraborty, et al. 2016) was used to merge the hybrid assembly with the PacBio only assembly, in which the latter was used as the reference assembly. However, assembly size of this merged assembly (NG50 =21.3Mbp; assembly size = 130Mbp) was similar to the hybrid assembly, and smaller than the estimated genome size of *D. melanogaster* females *(Hoskins, et al. 2002)*. Because the PacBio assembly size was closer to the estimated genome size, we added the contigs unique to the PacBio only assembly to the merged assembly using *quickmerge.* For this second round of merging (hco =5.0, c=1.5, l =5Mb), the merged assembly from the first round of merging was used as the reference assembly and the PacBio only assembly was used as the query assembly. Further improvements in assembly contiguity (N50 = 22.3Mb) was accomplished by running *finisherSC* (Lam, et al. 2015) with default settings on the final merged assembly. Next, the assembly was polished twice with *quiver* (as implemented in smrtanalysis v2.3) and once with *pilon* (Pilon 1.3) (Walker, et al. 2014). For pilon, we used the same Illumina reads (King, et al. 2012) that were used to generate the hybrid assembly.

### Bionano data

For collection of Bionano Irys data, A4 embryos of up to 12h of age were collected in apple juice-agar Petri dishes. Embryos were dechorinated using 50% bleach solution and passed through nitex nylon mesh to remove yeast and agar pieces. Approximately 250mg of embryo was placed into a prechilled Eppendorf tube and stored at -80C freezer. DNA was extracted following the manufacturer’s "Soft-Tissue” protocol (Bionano Genomics, San Diego). Frozen embryos were cut into <3mm pieces and placed into prechilled 500ul buffer HB per 10mg tissue and homogenized with 10 plunges in a Dounce/ Tenbroeck homogenizer. The homogenized tissue was incubated on ice for 5 minutes. 500 ul of the supernatant was transferred to a 1.5ml Eppendorf tube and an equal volume of ice cold ethanol was added to it. The ethanol was mixed by inverting the tube 10 times and then incubated on ice for 1 hour. The solution was centrifuged at 1500 *g at 4°C for 5 minutes and the supernantant was discarded. The pellets were resuspended in 66ul buffer HB and incubated at room temperature for 5 minutes. 40 ul prewarmed (43°C) low melting agarose was added to the buffer containing DNA, mixed with a pipette, and then solidified at 4°C. Five such agar plugs were transferred to a 50 ml tube and 2.5ml Lysis buffer and 200ul proteinase K was added to it. After an overnight incubation at 50°C for protein digestion, 50ul RNase A was added and incubated at 37°C for an hour to remove the RNA. The plugs were then washed 4 times with 10ml Wash buffer for 15 minutes at 180 rpm. Plugs are then transferred to a 1.5 ml tube with a spatula and melted at 70°C, followed by digestion of agarose with 2ul GELase at 43°C for 45 minutes. DNA was recovered by dialyzing DNA for 45 minutes on a membrane floating on 15ml TE at room temperature. The DNA was transferred to a 1.5 ml tube and quantified with Qubit BR assay kit.

The Bionano Irys optical data generated from the A4 DNA was generated and assembled with IrysSolve 2.1 at Bionano Genomics (San Diego, CA). The A4 Bionano assembly was then merged with the A4 assembly contigs with IrysSolve. To create the Bionano based scaffolds, assembly disagreements between the two were resolved by retaining the assembly features from the Bionano assembly where the two assemblies disagreed.

### Comparative scaffolding

The assemblies of all three genomes were scaffolded with a custom c++ program called *mscaffolder* (https://github.com/mahulchak/mscaffolder) using the release 6 *D. melanogaster* genome (r6.09) assembly (Hoskins, et al. 2015) as the reference. Prior to scaffolding, transposable elements and repeats in both assemblies were masked using default settings for *Repeatmasker (v4.0.6).* The repeatmasked A4 assembly was aligned to the repeatmasked major chromosome arms (X,2L,2R,3L,3R,4) of *D. melanogaster* ISO1 assembly using *MUMmer (Kurtz, et al. 2004).* Alignments were further filtered using the delta-filter utility with the -m option and the contigs were assigned to the specific chromosome arms based on the mutually best alignment. Contigs showing less than 40% of the total alignment for any chromosome arms could not be assigned a chromosomal location and therefore were not scaffolded. The mapped contigs were ordered based on the starting coordinate of their alignment that did not overlap with the preceding reference chromosome-contig alignment. Finally, the mapped contigs were joined with 100 Ns which represented assembly gaps. The unscaffolded sequences were named with a ‘U’ prefix.

### BUSCO analysis

To evaluate completeness and accuracy of the A4 assembly, *busco* (v1.22)(Simao, et al. 2015) was run on both scaffolded A4 assembly and the ISO1 release 6 assembly using the insect BUSCO database (total 2675 BUSCOs). Busco reported 5 BUSCOs (BUSCOaEOG75R3J9, BUSCOaEOG7SJRJ9, BUSCOaEOG7SJRK2, BUSCOaEOG7WMR0H, BUSCOaEOG71S8ZH) that are present in the ISO1 assembly as missing from the A4 assembly. To validate the absence of these 5 BUSCOs in the A4 assembly, all five genes (*Ftz-f1, CG7627, Raw, Maf1, Cv-c*) corresponding to the five BUSCOs were searched in the A4 assembly using full length sequence of the ISO1 genes (downloaded from FlyBase (dos Santos, et al. 2015)) using *MUMmer.* Surprisingly, the genome aligner *nucmer* found all five ‘missing BUSCOs’ to be present in the A4 assembly in single copies. Consequently, the BUSCO counts for A4 were adjusted accordingly.

### Structural variant detection

#### CNVs via whole genome alignment

To identify the copy number variants between iso1 and A4, we aligned the two genomes using *mummer* (Kurtz, et al. 2004) (mummer -mumreference -l 20 -b). The maximal exact matches (MEM) between the two genomes found by *mummer* were clustered using *mgaps* (*mgaps* -C -s 200 -f .12 -l 100). The l parameter in *mgaps* was set to 100 to detect duplicates that are 100bp or longer. We used a pipeline called *svmu* (Structural Variants from Mummer; https://github.com/mahulchak/svmu) to automate the copy number variants detection based on the overlapping *mgaps* clusters. When reference sequence regions in two separate alignment clusters overlapped, the overlapping segment of the reference sequence regions was inferred as duplicated in the query sequence. However, this can also potentially identify a duplicated sequence that is present in the both genomes but diverged due to the presence of repeats or indels around them. Furthermore, copy variants thus obtained also contain TE sequences, which were filtered using TE annotations by *Repeatmasker* (v4.0.6). False positives detected due to alignment issues were filtered by aligning the duplicated reference sequences back to the reference and A4 genomes using *nucmer (nucmer* -maxmatch -g 200) and then counting the copy number of each such sequence in each genome using *checkCNV,* which is also included in the svmu pipeline. The program *svmu* was run with the default parameters; *checkCNV* was run with c = 500 (max copy number 500), qco = 10000 (10kb of insertion/deletion allowed within a copy; this accounts for TE or other insertions of up to 10kb within a gene copy), rco = 0.2 (unaligned length of up to 20% of the sequence length between reference and query copies is allowed). CNVs (Table S9) that occurred 2kbp of each other were assumed to be part of a single mutation and therefore they were combined (using *bedtools* merge -d *2000*) *(Quinlan 2014)* for the purpose of counting total CNVs present in the genome. However, total sequence affected by CNVs was counted before merging was done. Functional annotation of the CNVs were made based on gene annotation of the release 6 of the reference genome.

#### Indels via whole genome alignment

Insertions (>100bp) in one genome is detected by looking for contiguous synteny in one genome that is broken by sequences that are longer than 100bp in the other genome. To find insertions in the A4 genome, we aligned ISO1 (reference) and A4 (query) chromosome arms using *nucmer* (default parameters). Next, we looked for alignment gaps wherein two adjacent syntenic segments in A4 are separated by more than 100 bp whereas the same adjacent syntenic segments in ISO1 are separated by less than 10% of insert length in A4. Indels are detected by a custom c++ utility called *findInDel* which is also part of the svmu pipeline (https://github.com/mahulchak/svmu).

#### Inversions via whole genome alignment

To identify the inversions in the A4 genome, the A4 genome was aligned to the ISO1 genome using *nucmer* (-mumreference). The delta file was converted into a tab delimited file called “aln_summary.tsv” using *findInDel.* The query (A4) genomic ranges that ran in the reverse direction with respect to the reference (ISO1) were recorded as inversions. TEs were removed from this list using a *Repeatmasker* annotated TE list for ISO1.

#### Genotyping CNVs, indels, and inversions using Illumina reads

Three common strategies are typically employed to discover copy number variants using Illumina high throughput short reads. One strategy uses variation in mapped read depth as the signal for presence of copy number variation, another uses orientation anomalies of paired end reads as signals for duplication, and the third strategy uses the mapping properties of split reads to discover the presence of structural variation breakpoints (Alkan, et al. 2011a). We used all three of the strategies because they exploit complementary aspects of the data (Alkan, et al. 2011a). We used *CNVnator* (Abyzov, et al. 2011) for read depth, *pecnv* for read pair orientation (Rogers, et al. 2014), and *pindel* (Ye, et al. 2009) for split read mapping approaches of duplicate discovery. We used 70X paired end A4 reads (King, et al. 2012) for finding duplicates in the A4 strain. Briefly, the reads were mapped to the release 6 reference sequence using *bwa mem* for *CNVnator* and *pindel* and *bwa aln* for *pecnv (Li and Durbin 2009).* The sam files containing the alignments were converted to bam files and sorted using *samtools (Li, et al. 2009).* The sorted bam files were used for CNV calling. For pecnv, we used a coverage cutoff of 3 following (Rogers, et al. 2014). To filter out the false positives. For *CNVnator,* we used a bin size 100 due to the high coverage of the data. Furthermore, we restricted our analysis on genotype comparison to CNVs that are 100bp or long and 25Kb or shorter. To genotype the large indels (>100bp) using Illumina data, we used *CNVnator* and *Pindel* using the same command line settings as used for the CNV calls. For inversion genotyping, *Pindel* was used.

TE insertion coordinates for A4 were obtained from flyrils.org (Cridland, et al. 2013). We restricted comparison of our TE insertion calls and that from (Cridland, et al. 2013) to the chromosome arm 2L because the genotypes are based on ISO1 release 5 coordinates and only chromosome 2L coordinates have remained unchanged between release 5 and release 6 (the assembly version used here). Furthermore, a single chromosome arm contained <150 indel mutations, which facilitated manual validation of each mutation from long read alignment. For similar reasons, manual validation CNV calls across different Illumina based methods was done for chromosome arm 2L.

### SNP and small indel detection

SNPs and small indels (<100bp) in the A4 assembly were identified using the *show-snps* utility from the *MUMmer* package(Kurtz, et al. 2004). First, A4 scaffolds were aligned to the ISO1 scaffolds using *nucmer (-mumreference)* To minimize spurious SNP calls due to repeats, repeats were filtered using *delta-filter* in conjunction with the –r amd –q options. SNPS and small indels were called from the filtered delta file using *show-snps* (using –Clr options).

### Validation of duplicates and indels

All duplicate and indel calls were examined by inspecting the dot plots of the duplicated and inserted-deleted sequences. Furthermore, to rule out assembly errors as the source of indel or CNV calls, we mapped the long reads to the A4 and ISO1 assembly using *blasr* v1.3.1.142244 (-bestn 1 –sam). The sam files were converted into bam files using *samtools view* command (samtools 1.3) and then sorted the bam files using *samtools sort* with the defaults parameters (Chaisson and Tesler 2012; Li, et al. 2009). We tested all CNVs and indels present in the chromosome 2L and examined the mapped reads at the genomic regions containing the inferred CNVs or indels using Integrative Genomics Viewer (IGV) (Thorvaldsdottir, et al. 2013). Furthermore, duplication of all full length genes was examined and validated using the mapped reads.

### Expression analysis

Genomewide gene expression difference between A3 and A4 larvae were analyzed following the method of (Marriage, et al. 2014). Sequences of the A3 genes were obtained from a A3 genome assembly constructed with publicly available A3 Illumina paired end reads(King, et al. 2012).To compare the gene expression level of the *Cyp28d1, CG7742,* and *Ugt86Dh* gene copies, we aligned the publicly available 100bp single ended RNAseq reads (Marriage, et al. 2014) to the A4 mRNA sequences using *bowtie2* (Langmead and Salzberg 2012) with the parameter--score-min L,0,0 to ensure that only perfect alignments (cigar string =100M) were retained. Only perfectly aligned reads were kept for downstream analysis so that reads specific to a gene copy could be obtained. We counted the unique perfectly aligned reads for each paralog and then calculated FPKM from these. Total number of reads aligned to the genomes were calculated based on the alignment of the single ended RNAseq reads aligned to the A4 and A3 genomes using tophat (Trapnell, et al. 2012). Because only reads overlapping the SNPs were counted for FPKM calculation, the transcript length was adjusted by subtracting the transcript length to which no SNP covering read aligned. That is reads aligning within 99bp of a SNP was not counted for FPKM calculation. For example, the *Cyp28d1* gene copies are distinguishable by 15 snps so when only perfectly aligned unique reads are counted, the effective transcript length of the *Cyp28d1* gene copies used in calculation of FPKM becomes 1509-(310) = 1199bp. Similarly, for *Ugt86Dh* and CG7742, transcript lengths of 1065 bp and 755bp were used to calculate FPKM, respectively. No such adjustments were made for the single copy genes.

### Testing for selective sweeps

Testing for natural selection was performed using the composite likelihood ratio (CLR) statistic for a recent selective sweep (Nielsen, et al. 2005), computed using the *SweepFinder2* software (version 1.0)(DeGiorgio, et al. 2016). CLR values were calculated using the frequency of SNPs present in each sample over a grid with 250 bp increments. Sites were polarized using an outgroup of three closely related species, where the ancestral state was inferred by sites that shared the same genotype across the release 2 reference genomes of *D. simulans, D. yakuba,* and *D. erecta.* Invariant sites that differed from the inferred ancestral state (substitutions) were included in the analysis, thus improving power and robustness to bottlenecks (Huber, et al. 2016; Nielsen, et al. 2005). The significance of the results was evaluated by comparing the CLR values to coalescent neutral simulations generated using the software *ms* (Hudson 2002).

### Estimating duplicate allele frequencies

The frequency of duplicate alleles was estimated from next-generation Illumina data by analyzing the density of divergently mapped read pairs. Reads were mapped against the release 6 ISO-1 reference genome using *bwa mem* (Li and Durbin 2009). Divergent read pairs were selected by taking the complement of paired reads in the BAM file that mapped with proper orientation, defined as pairs of reads that mapped to the same chromosome on opposite strands and were flagged by the aligner as being properly aligned with respect to the each other. Duplications were called for samples that showed a clear peak and high signal-to-noise ratio in the coverage density for divergent read pairs at breakpoints surrounding genes that were found to be duplicated in A4. The divergent read pair signals for several duplicate alleles for *Cyp28d1* from various populations are shown in Figure S24.

## Supplementary materials

**Figure S1.**
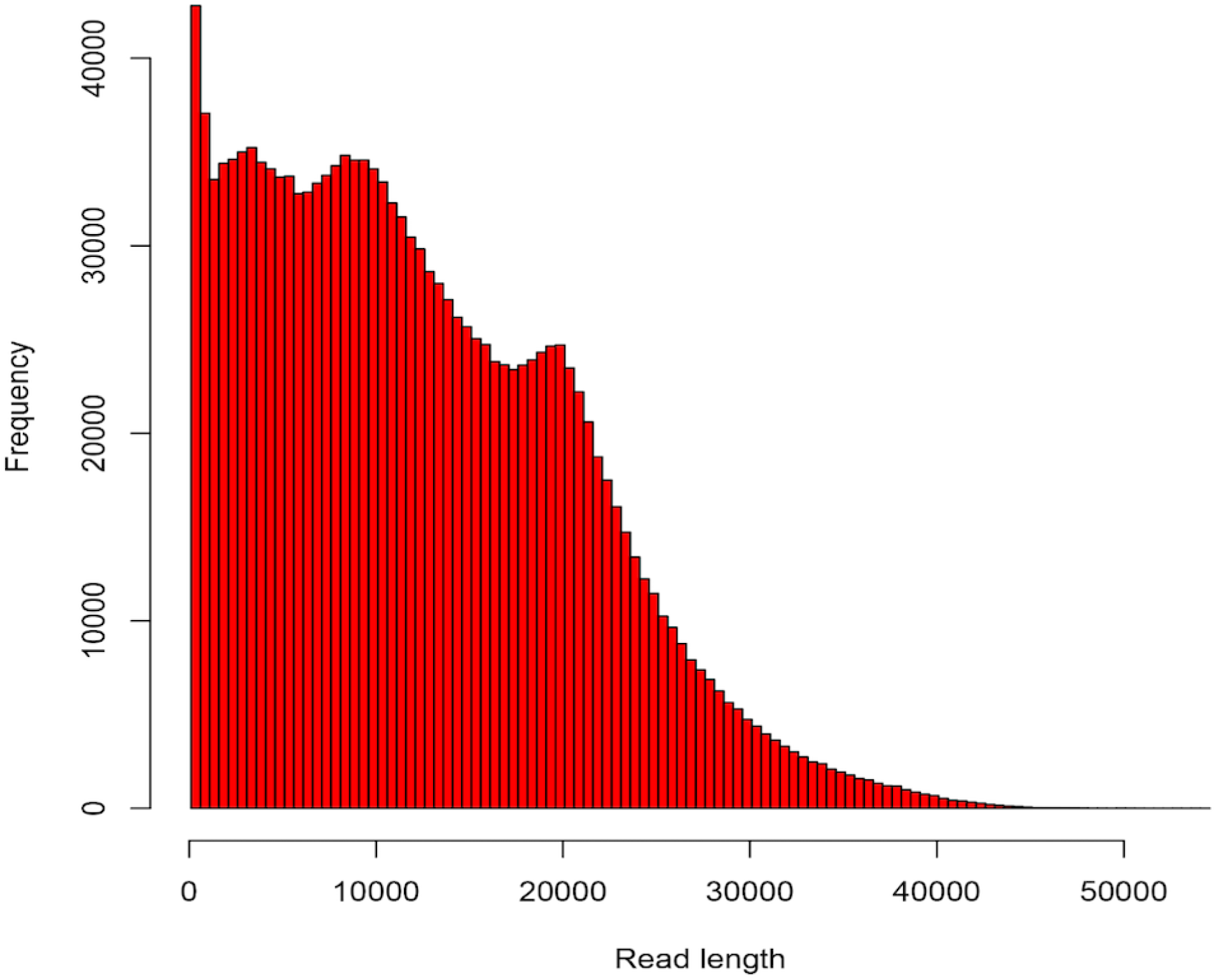
Distribution of A4 long reads. 50% of total coverage is contained within read length 18 kbp or longer (ie NR50 = 18 kbp).

**Figure S2.**
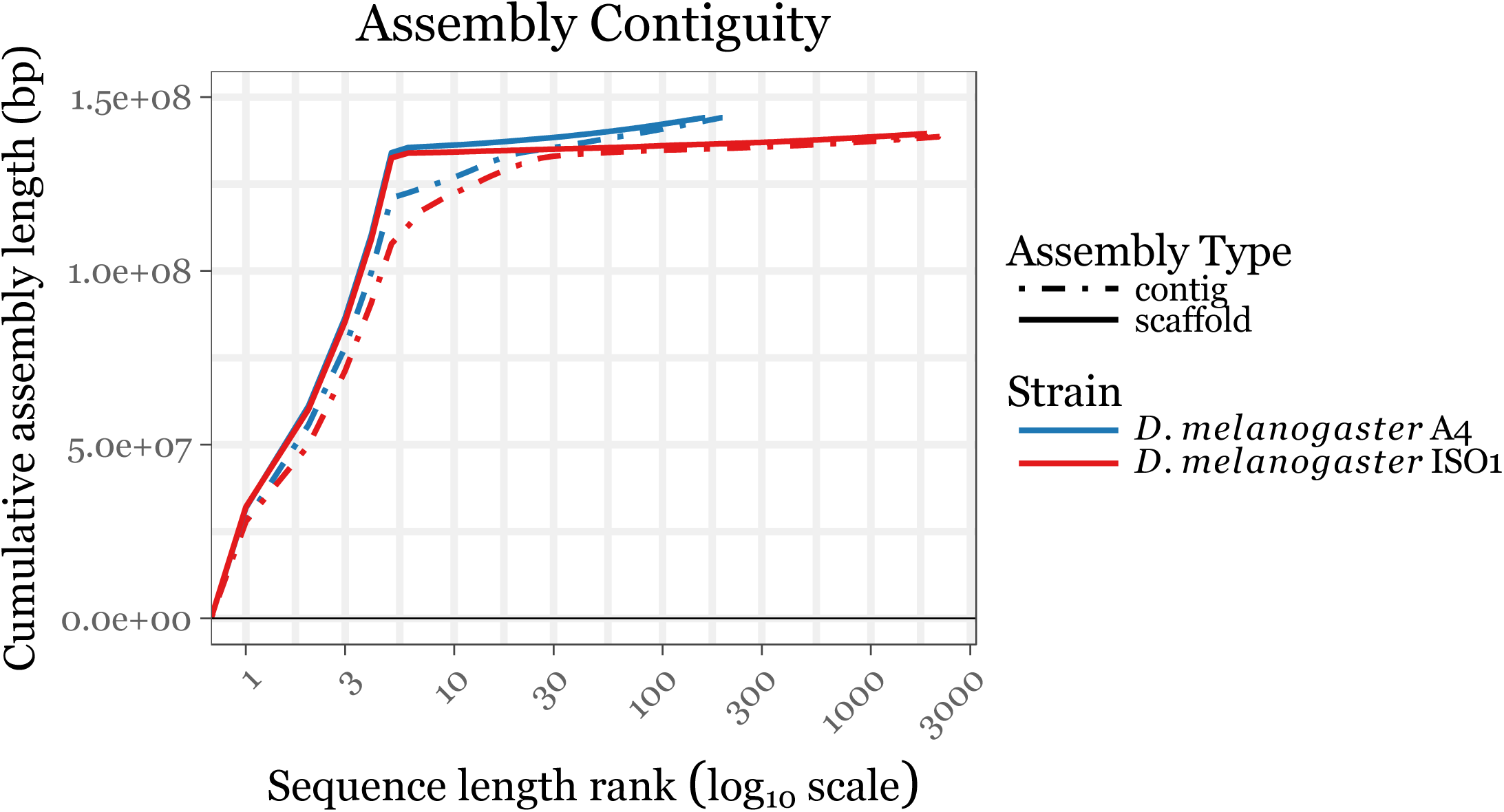
Cumulative sequence length distributions for the A4 and ISO1 assemblies. The X-axis is the sequence length rank sorted in descending order (i.e. the largest sequence is rank 1, the 2^nd^ largest is rank 2, etc.) The Y-axis is the cumulative length of all sequences to the rank on the X-axis. A4 is more contiguous than ISO1 on both the contig and scaffold levels. The total amount of genome assembled for A4 is also about 4 Mbp more than for ISO1, as indicated by the A4 curves reaching a higher Y-value. The ISO1 assemblies are derived from the release 6 version from <u>FlyBase.</u> The mitochondrial genome was excluded from both genomes and the Y-chromosome sequences were excluded from the ISO1 genome (the A4 genome has no Y-chromosome sequence, as it was assembled from reads derived from females).

**Figure S3.**
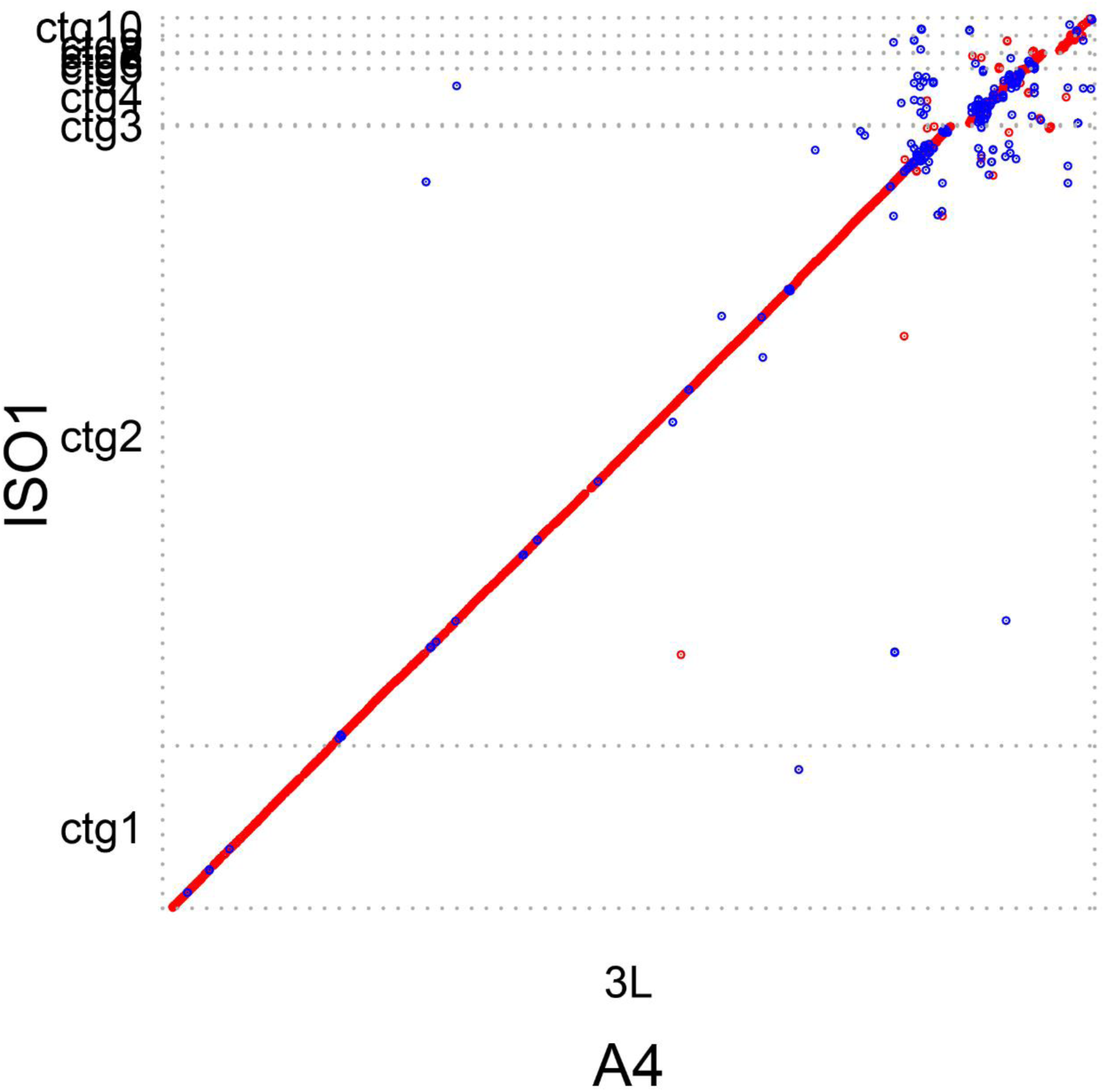
Alignment dot plot between the 3L contigs in ISO1 and a single 28Mb contig in A4 showing higher assembly contiguity in A4. As evidenced here, the A4 3L has fewer gaps than the reference 3L assembly.

**Figure S4.**
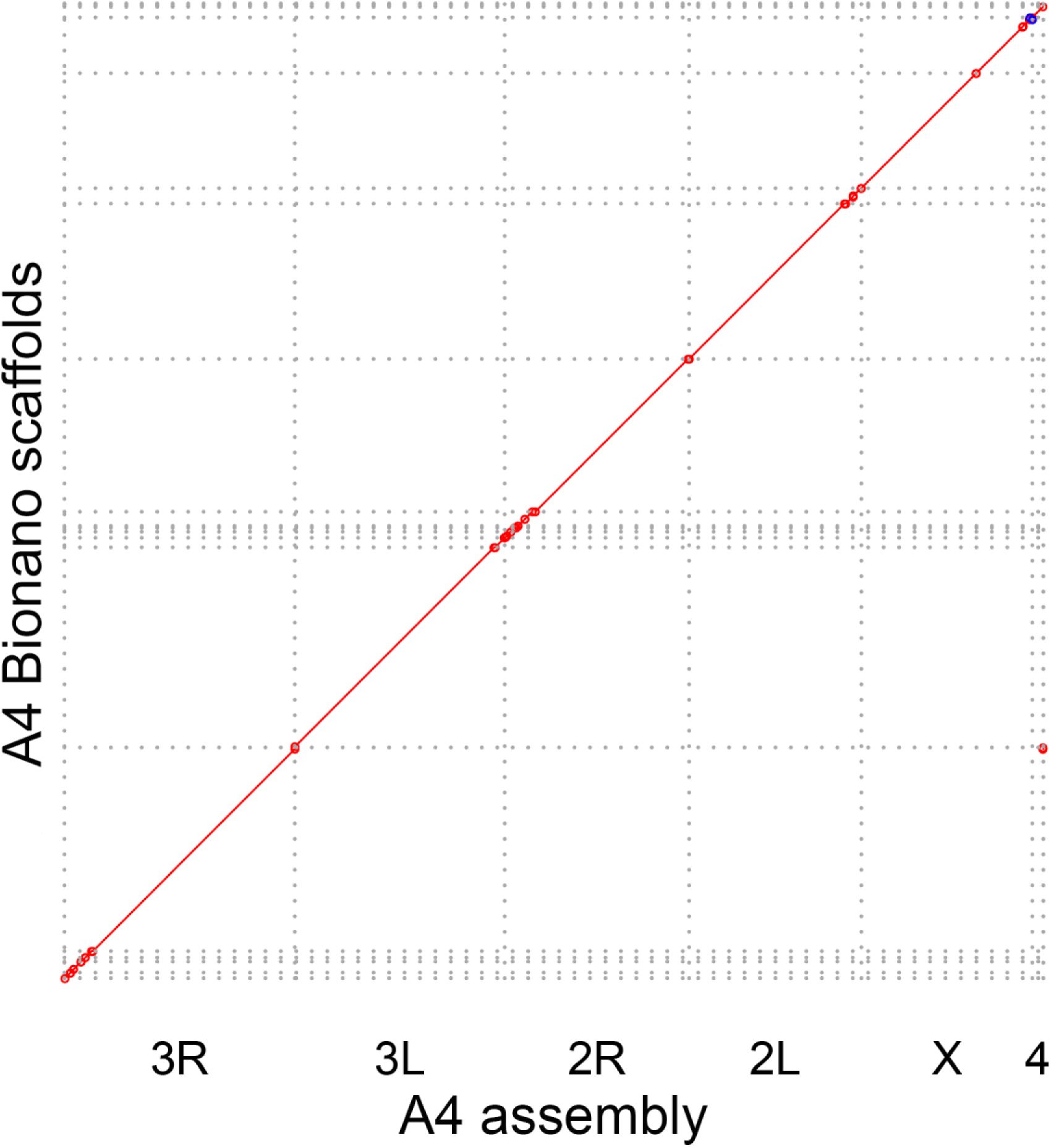
Alignment dot plot between the A4 assembly scaffolded using the reference assembly (Hoskins, et al. 2015) and an assembly scaffolded with a Bionano optical map. Collinearity of the two assemblies suggest that the contiguity of A4 assembly is not a result of incorrect contig joining. Evidence of a strain specific inversion (blue) mapping to the distal end of the X chromosome is present in the Bionano assembly, but absent in the PacBio assembly.

**Figure S5.**
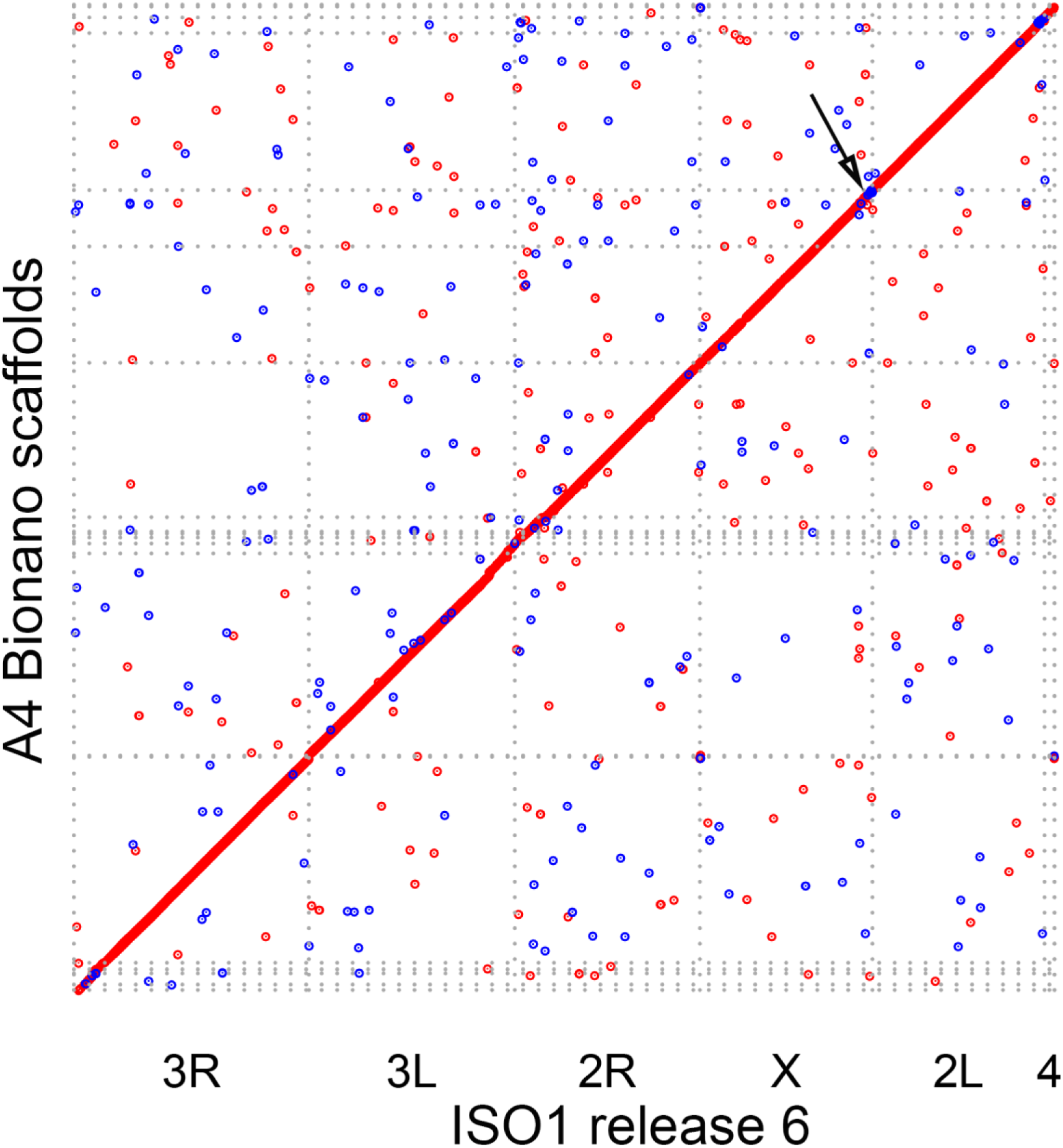
Alignment dot plot between the ISO1 release 6 scaffolds and the A4 Bionano assembly scaffolds. The small off-diagonal alignments (dots) are due to TE and repeats. The evidence of the X chromosome inversion (arrow) at the pericentric heterochromatin of A4 X chromosome is visible in a Bionano scaffold mapping to the distal end of ISO1 X.

**Figure S6.**
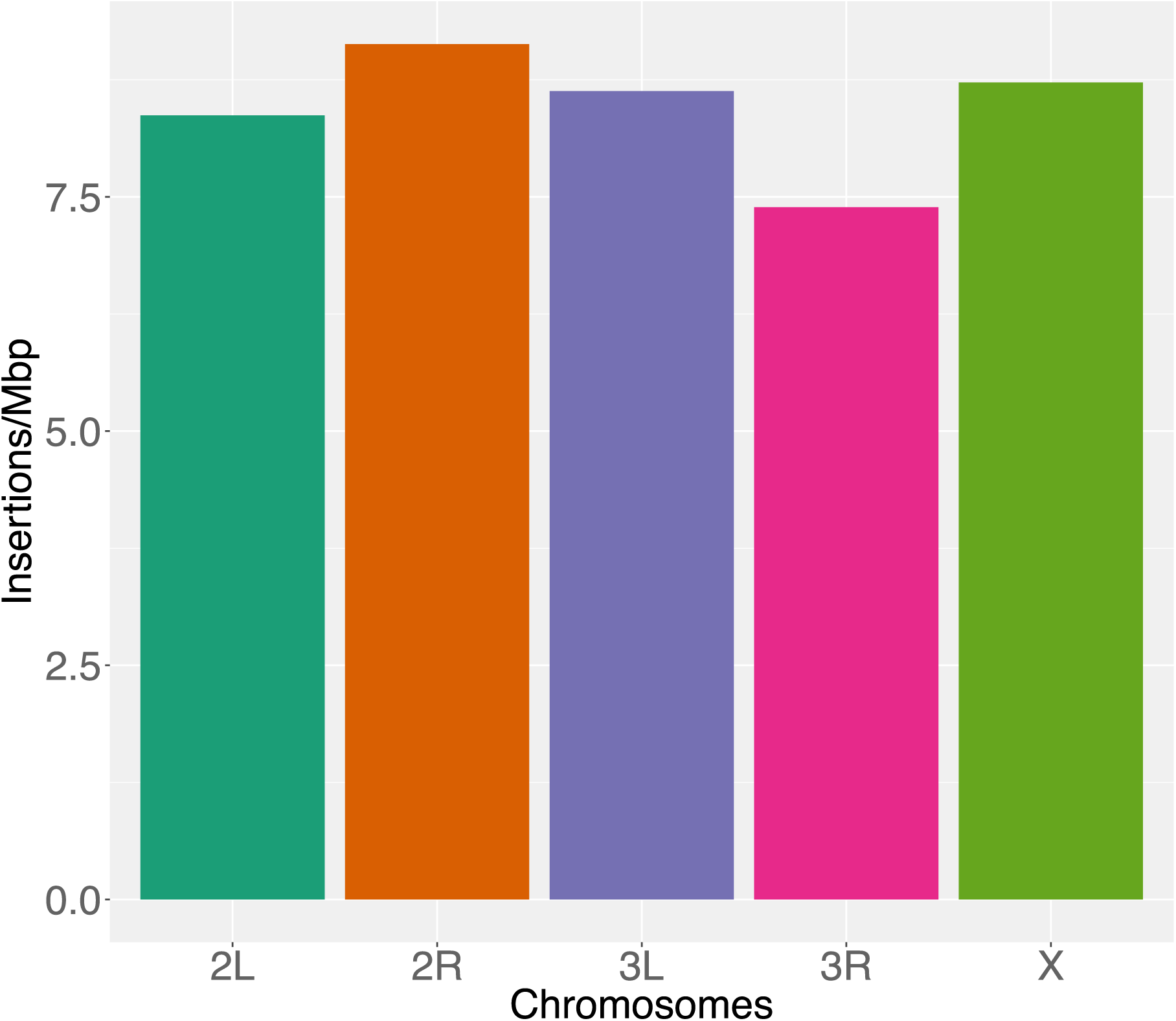
Number of insertions per megabase of euchromatic DNA in each chromosome arm

**Figure S7.**
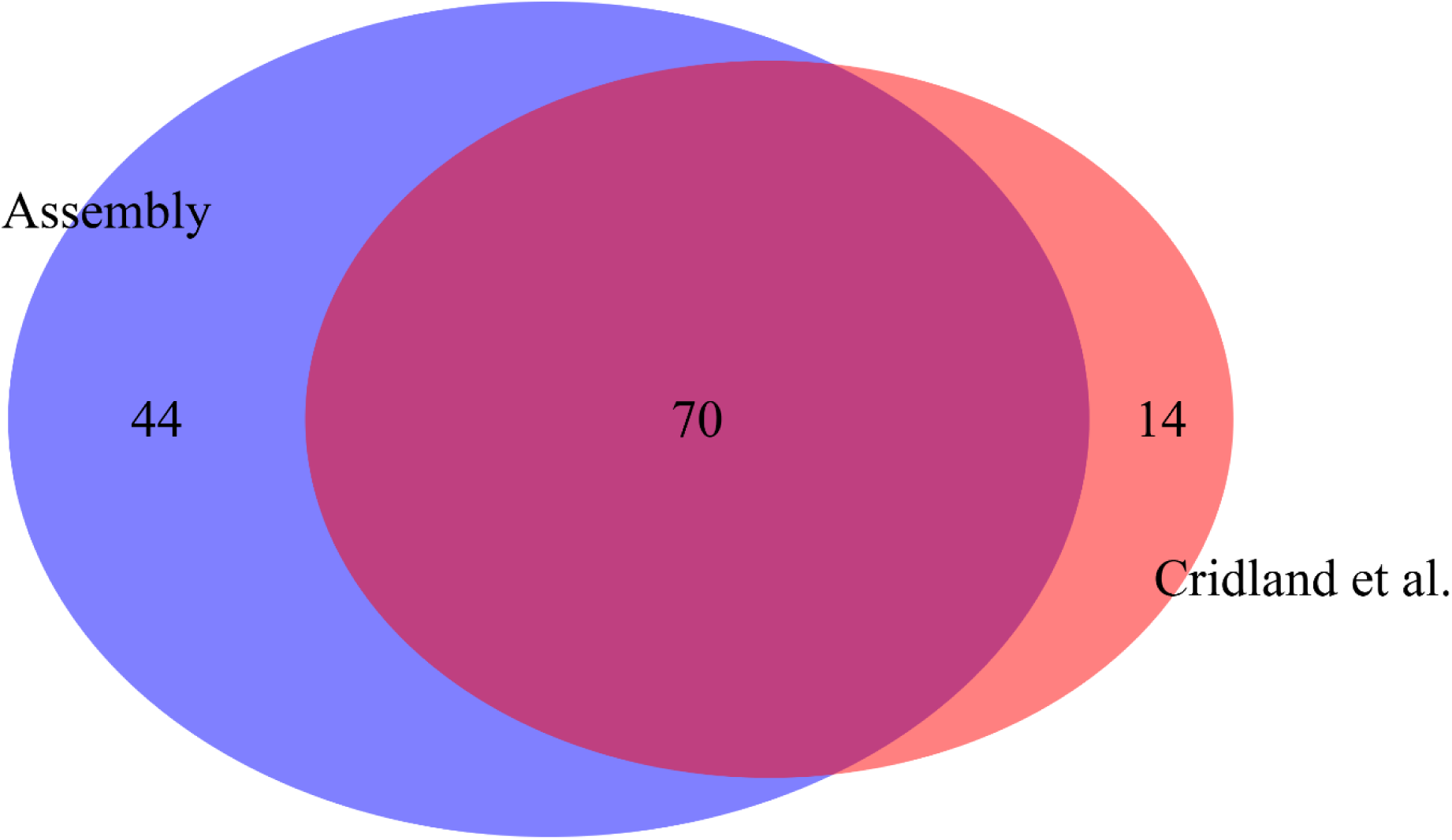
The short-read based TE insertion genotyping method of (Cridland, et al. 2013) can detect 63% of the TEs insertions present in the euchromatic regions of the 2L chromosome arm of A4 assembly. All TE calls for all categories above were manually curated. The 37% of TEs missed are considered hidden/invisible to short-read methods.

**Figure S8.**
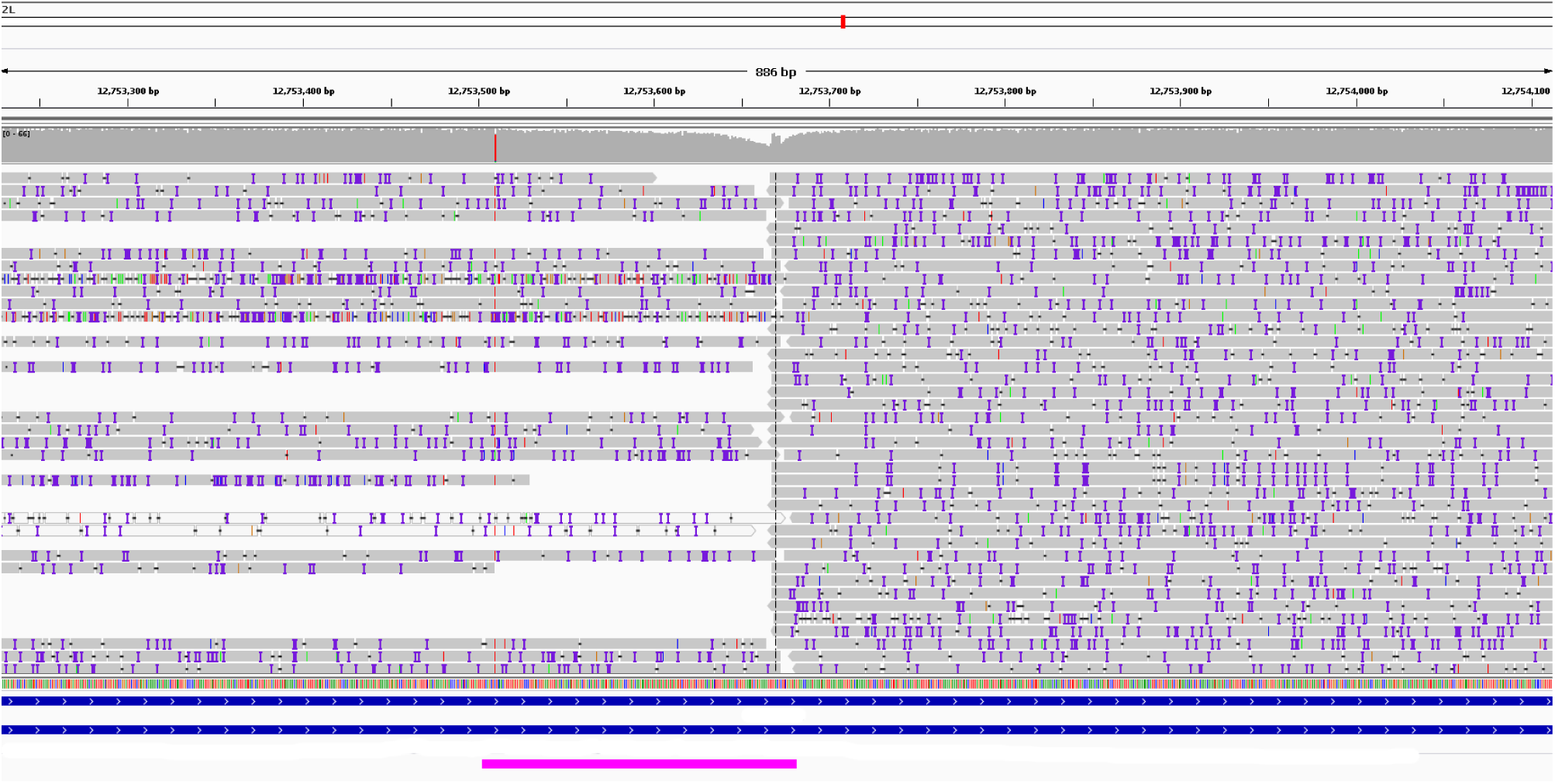
A TE insertion that is absent in the list of A4 TE insertion genotypes based on paired end Illumina reads(Cridland, et al. 2013; Cridland and Thornton 2010). The insertion point, indicated by the drop in A4 long read coverage (aligned to the release 6 ISO1 assembly), is located right next to an existing TE called INE-1(Marriage, et al. 2014)2210 (shown in pink) inside an intron of the gene *MRP* (Multidrug-Resistance like Protein 1). Such existing TEs obscure the TE insertion signals that short-read based method(Cridland and Thornton 2010) uses to detect TE insertions.

**Figure S9.**
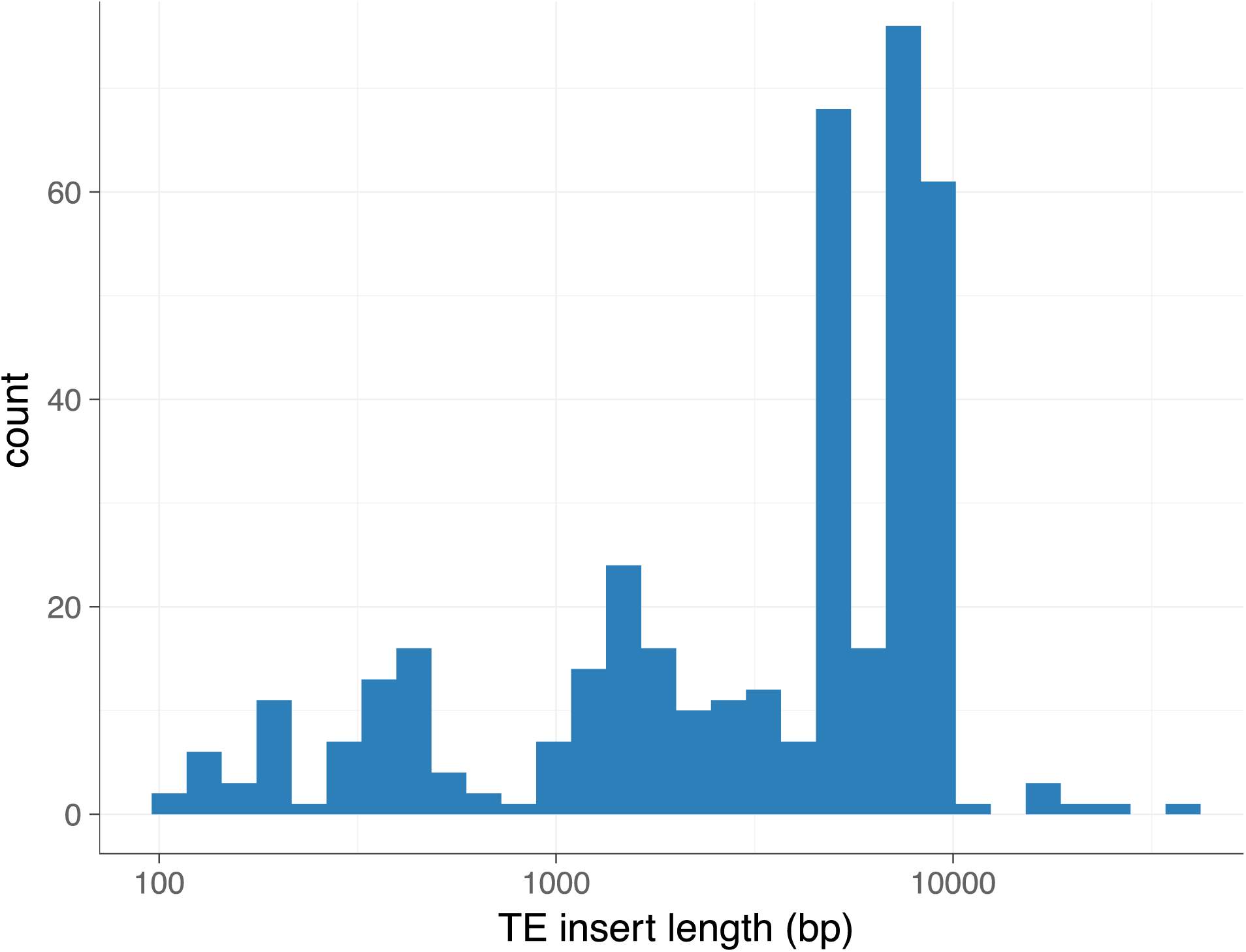
The distribution of lengths of the TEs that insert within introns. More than 50% TEs inserting within introns are large (>5kbp; median 5016bp) and may cause ‘intron delay’(Swinburne and Silver 2008).

**Figure S10.**
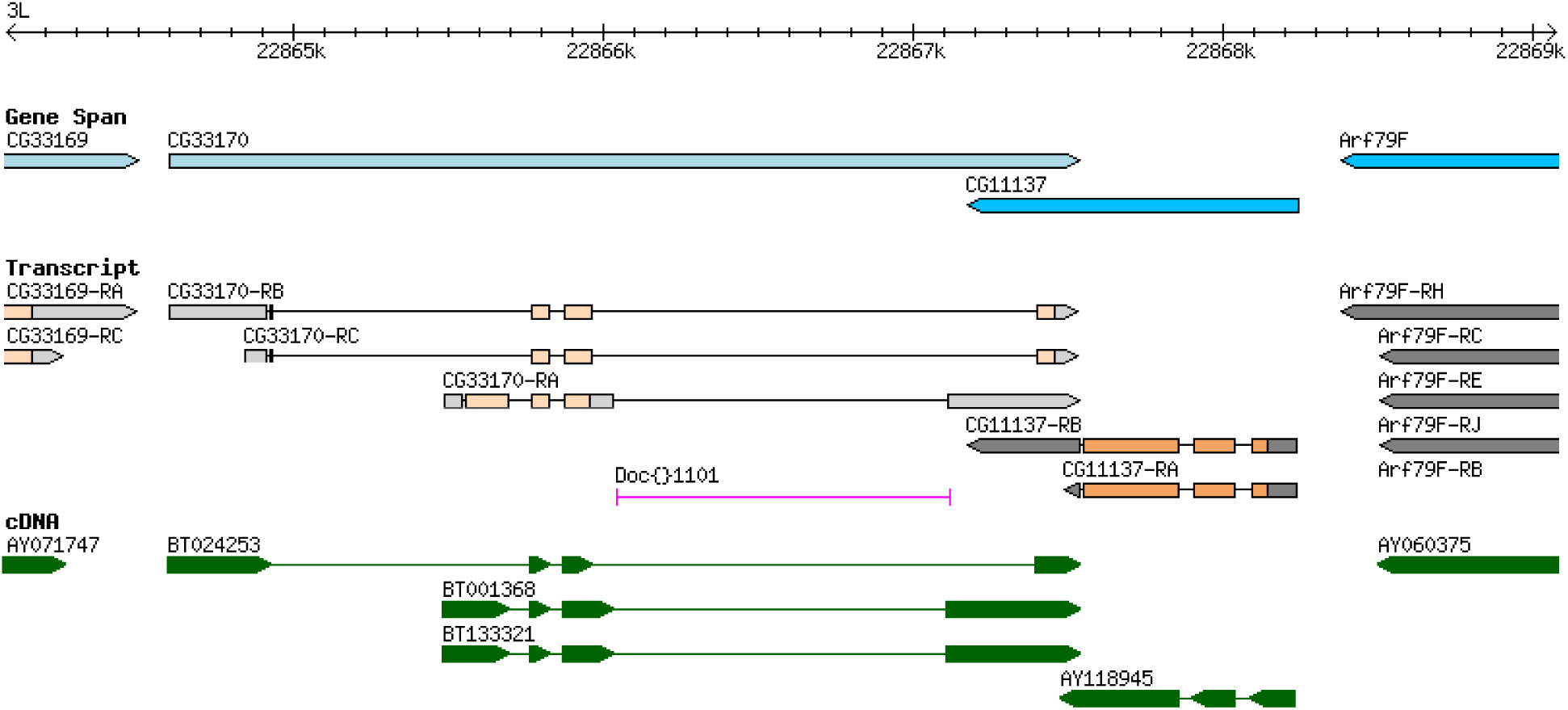
Insertion of a private TE (*Doc* element) in the ISO1 gene *CG33170* corresponds to the whole 4^th^ intron in one of the ISO1 *CG11137* isoforms as shown in FlyBase (dos Santos, et al. 2015) annotation of the gene. The TE is private to ISO1.The cDNA annotations are based on Drosophila Gene Collection(Stapleton, et al. 2002).

**Figure S11.**
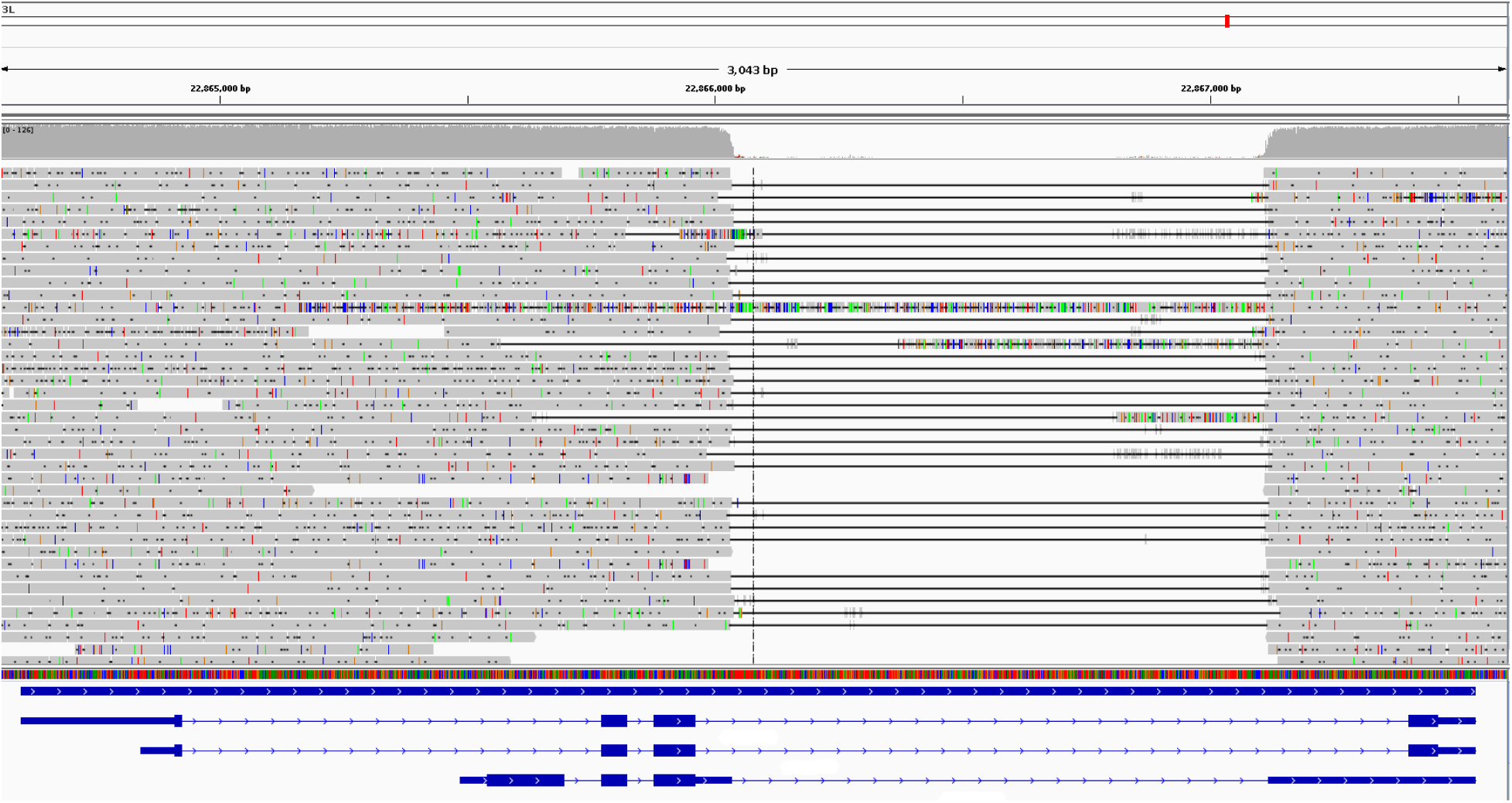
The A4 long reads aligned to ISO1 release 6 assembly to show the ISO1 specific *Doc* element insertion (supplementary Fig. S10) in the gene *CG33170.* As demonstrated by the schematic diagrams (blue lines and rectangles) of the *CG33170* transcripts, this private TE in ISO1 covers the entire intron in one of the transcript isoforms of *CG33170.*

**Figure S12.**
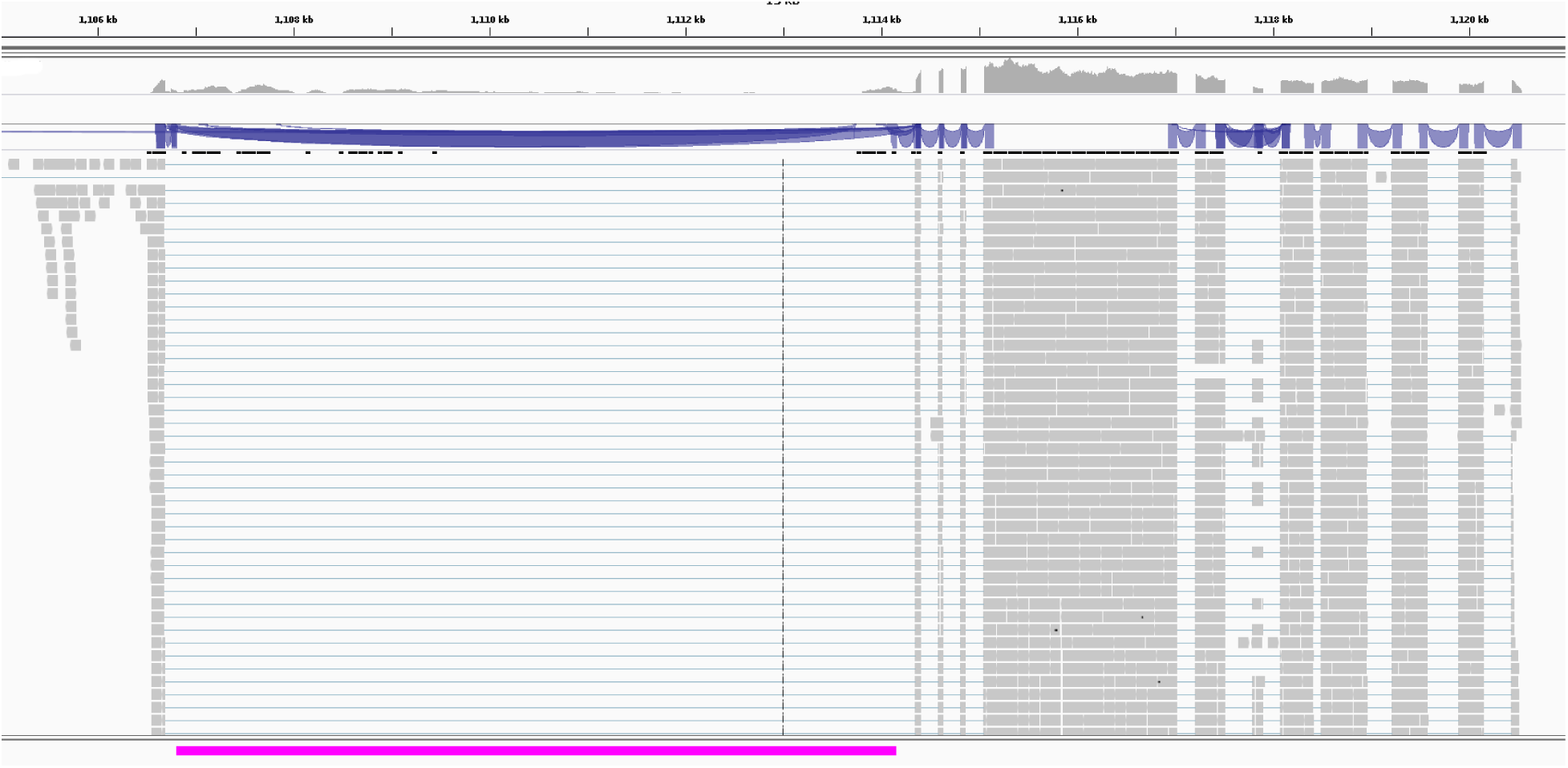
Example of a new intron in *CG13900* of A4 gained via MDG1 LTR insertion (the pink bar). A4 RNAseq reads(Marriage, et al. 2014) were mapped to the A4 assembly using Tophat. The blue lines and the purple ribbons indicate intron and splice junctions. For this TE intron, individual reads span the entire 7348bp insertion suggesting that the TE is spliced out. Grey rectangles indicate reads/exons.

**Figure S13.**
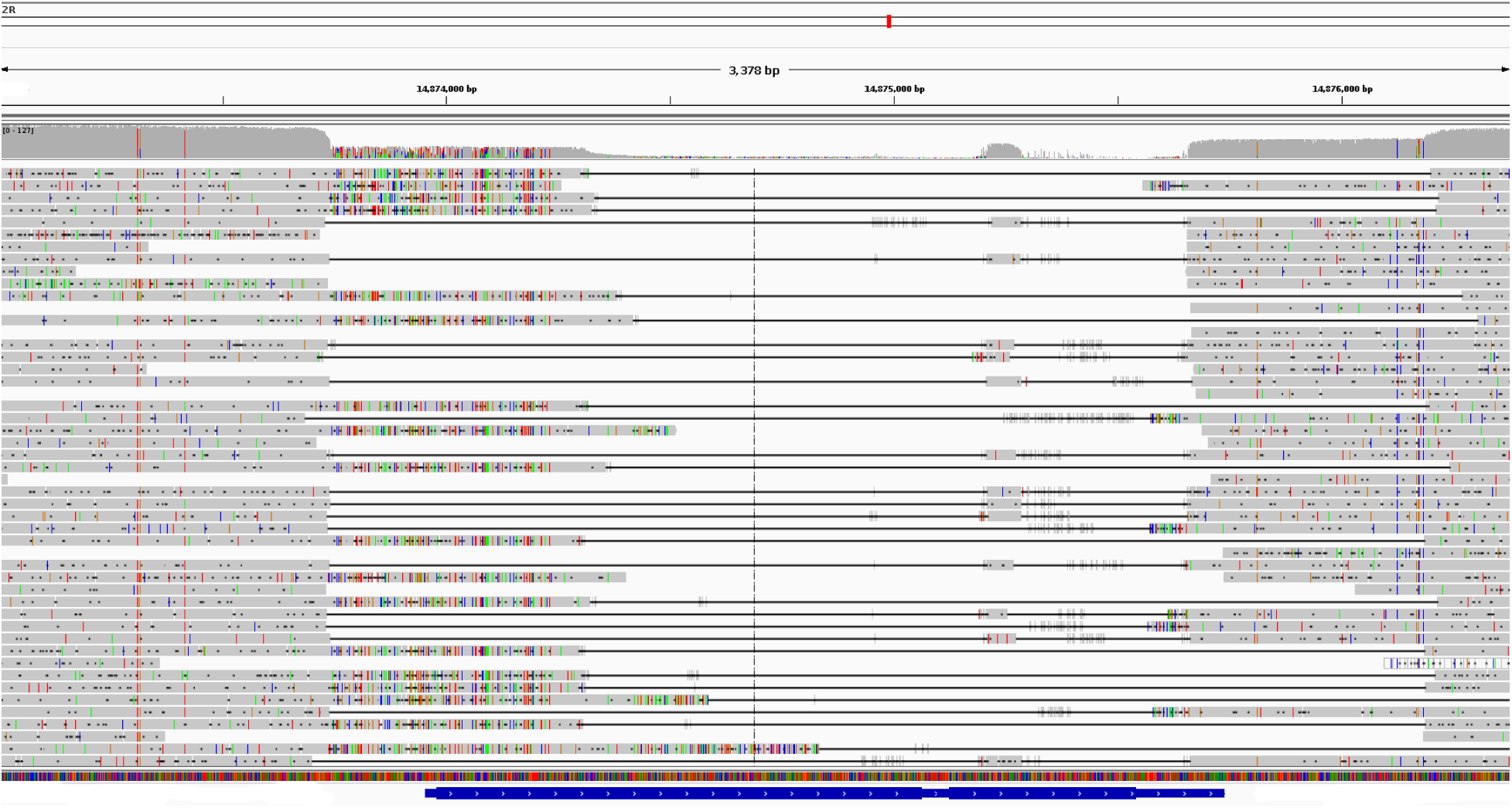
A4 read coverage at the ISO1 genomic region containing *Cyp6a17.* The ISO1 *Cyp6a17* gene model is shown in dark blue. As evident here, A4 is lacking *Cyp6a17* and might show defects in temperature preference(Kang, et al. 2011).

**Figure S14.**
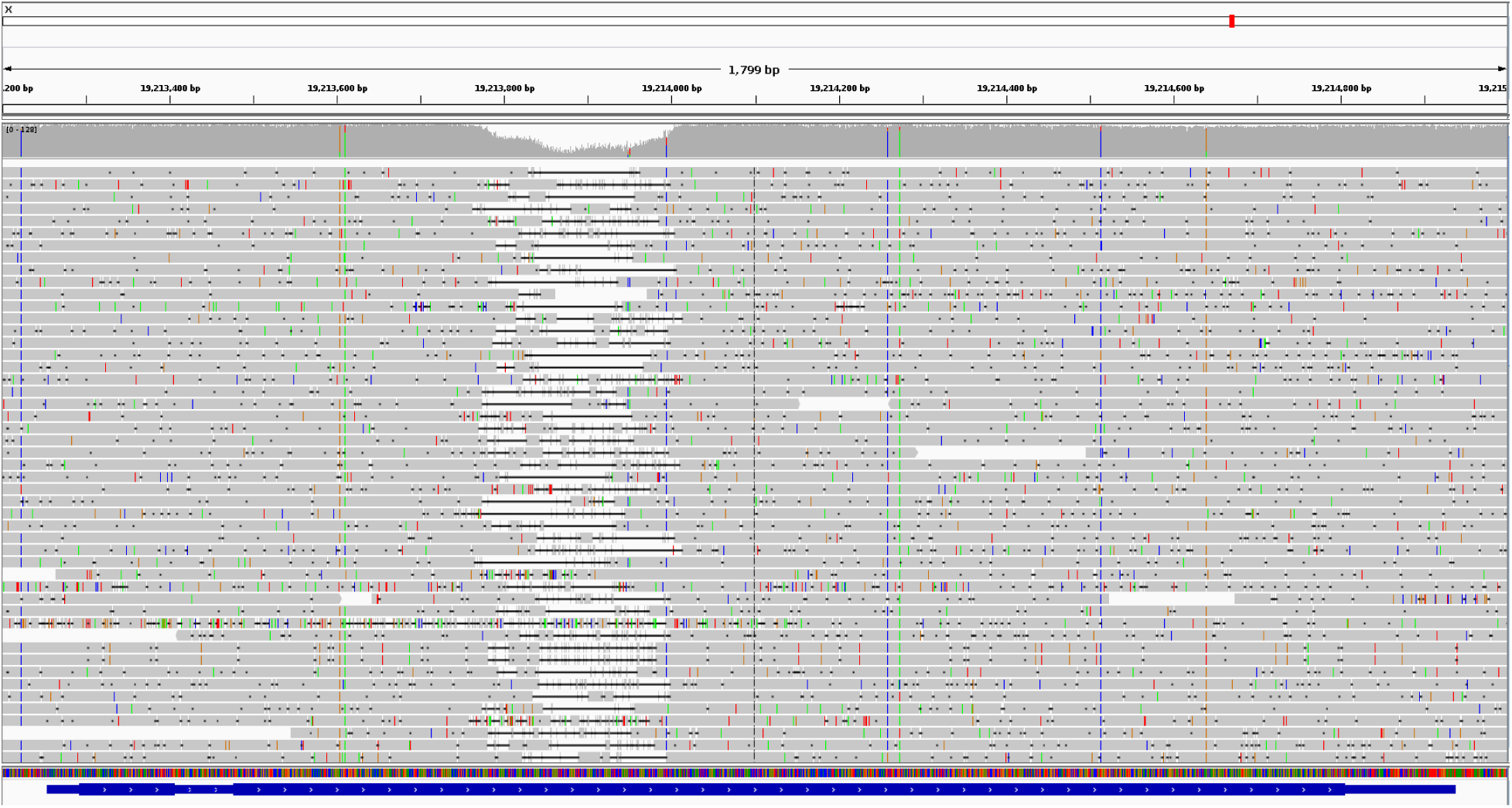
Deletion in *Mur18B* gene in A4 as shown by the presence of an alignment gap between the A4 long reads and genomic region the X:19213350-19214900 in the ISO1 release 6 assembly. The deletion removes 129bp from the ORF of the gene.

**Figure S15.**
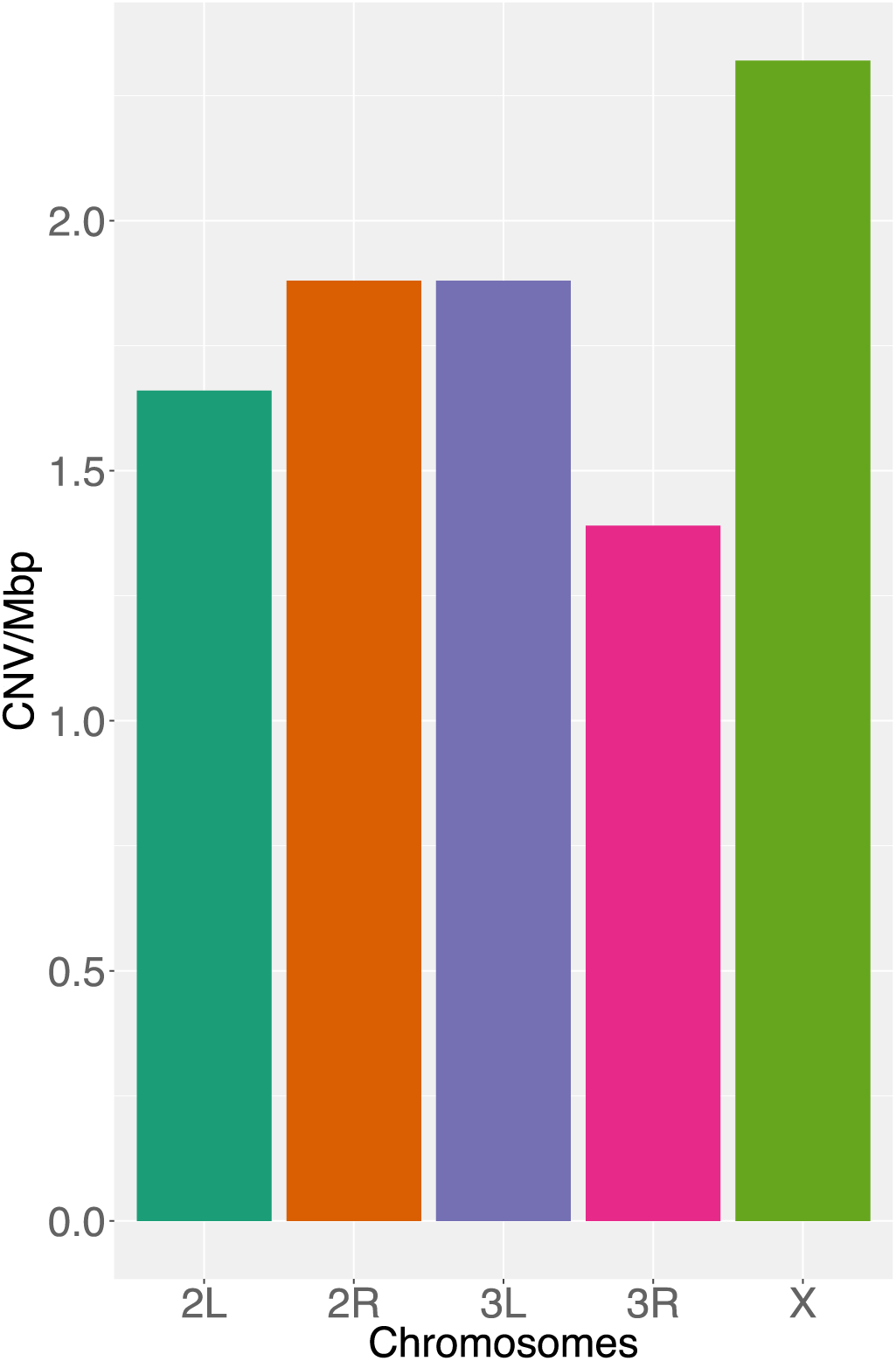
Number of duplicates per megabase of euchromatin.

**Figure S16.**
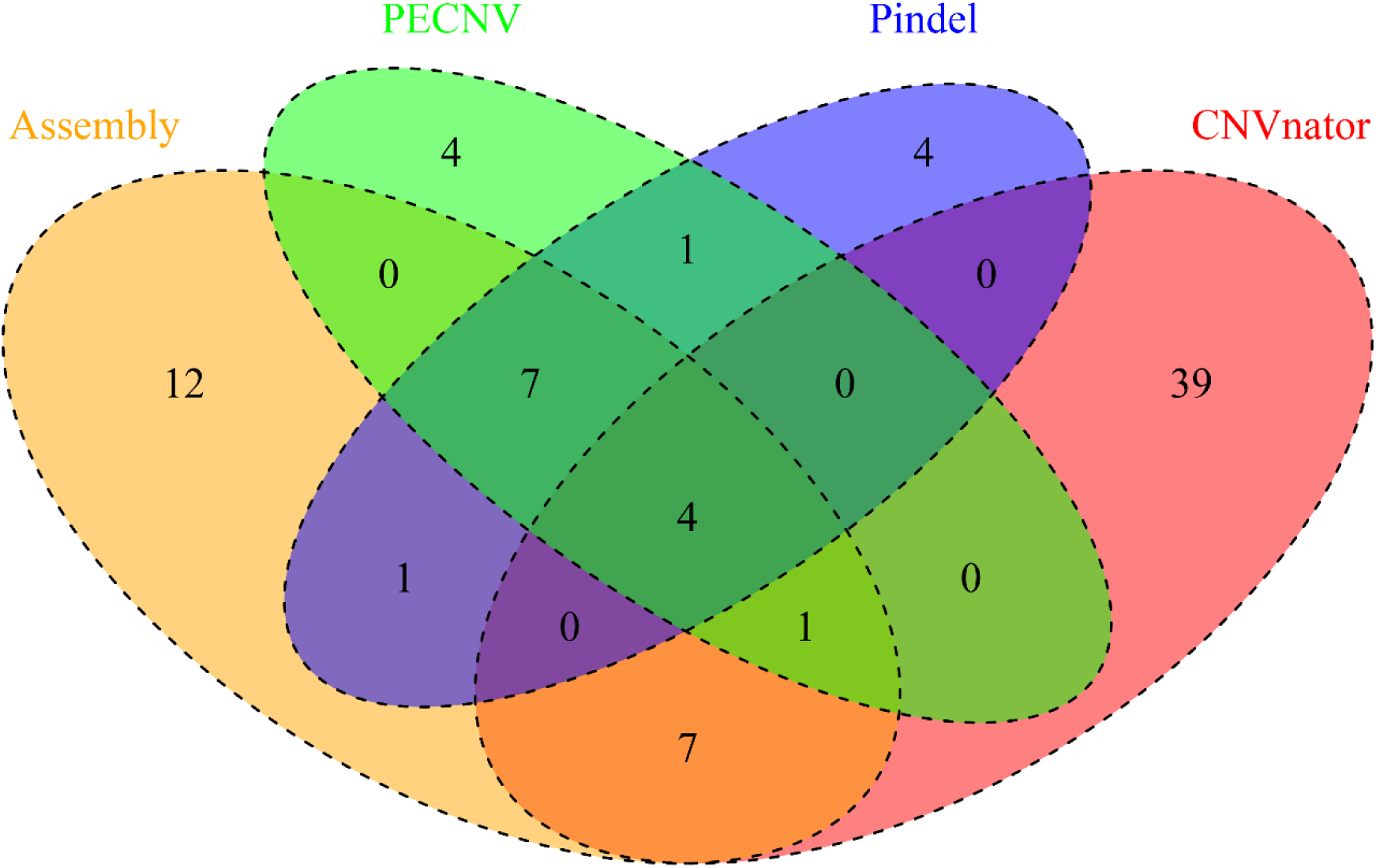
Tandem duplications detected by our A4-ISO1 genome alignment approach (Assembly), a split read mapping method (*Pindel* (Ye, et al. 2009)), a read pair orientation method (*PECNV* (Rogers, et al. 2014)), and a read coverage based method (*CNVnator* (Abyzov, et al. 2011)) on chromosome arm 2L. All mutation calls were manually curated. Mutations detected by *PECNV* and *Pindel* overlap significantly, whereas CNVs detected by *CNVnator* overlap little with those detected by *Pindel* and *PECNV. CNVnator* calls, when not confirmed by other methods, are dominated by false positives and yield a similar false negative rate as the other methods. When used in concert with *PECNV,* it permits discovery of one additional mutation on 2L. Mutations discovered by any two methods were considered discoverable by short reads. Other mutations are considered hidden/invisible.

**Figure S17.**
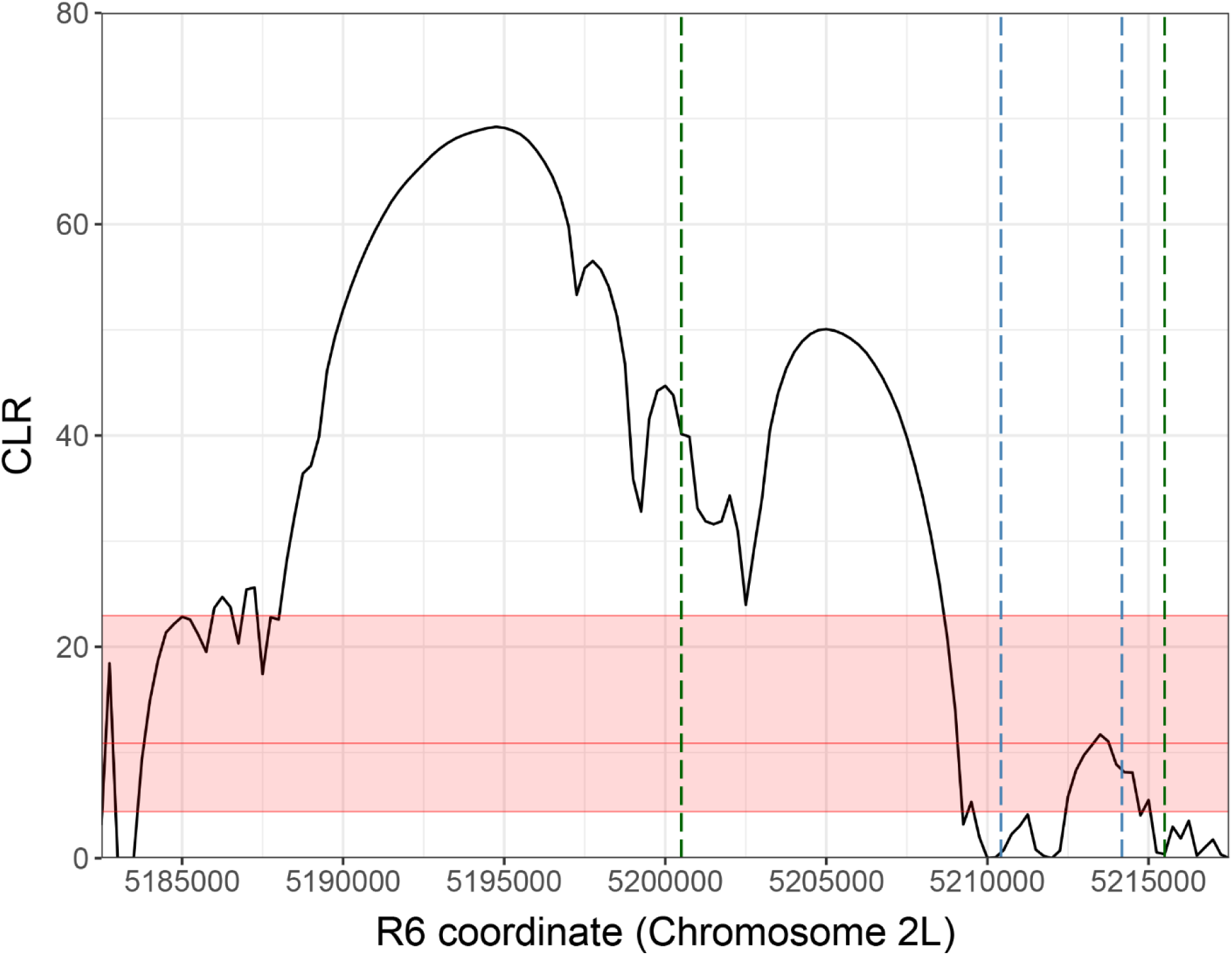
Distribution of composite likelihood ratio statistic (Nielsen, et al. 2005) of single nucleotide site frequency spectrum at the genomic region containing *Cyp28d1* in the French population from (Bergman and Haddrill 2015). The CLR peak falls immediately adjacent to the duplication. Given problems genotyping SNPs in duplicates, we do not actually expect the paralogous regions to be easily interpretable in a CLR framework. The red shaded region represents the empirical 95% confidence interval for the maximum CLR values based on 100 neutral simulations using the observed SNPs at this region (Materials and Methods). Vertical green lines indicate the span of all duplication alleles observed in all populations surveyed in Figure 3a. Vertical blue lines indicate the extent of the duplication discovered in A4.

**Figure S18.**
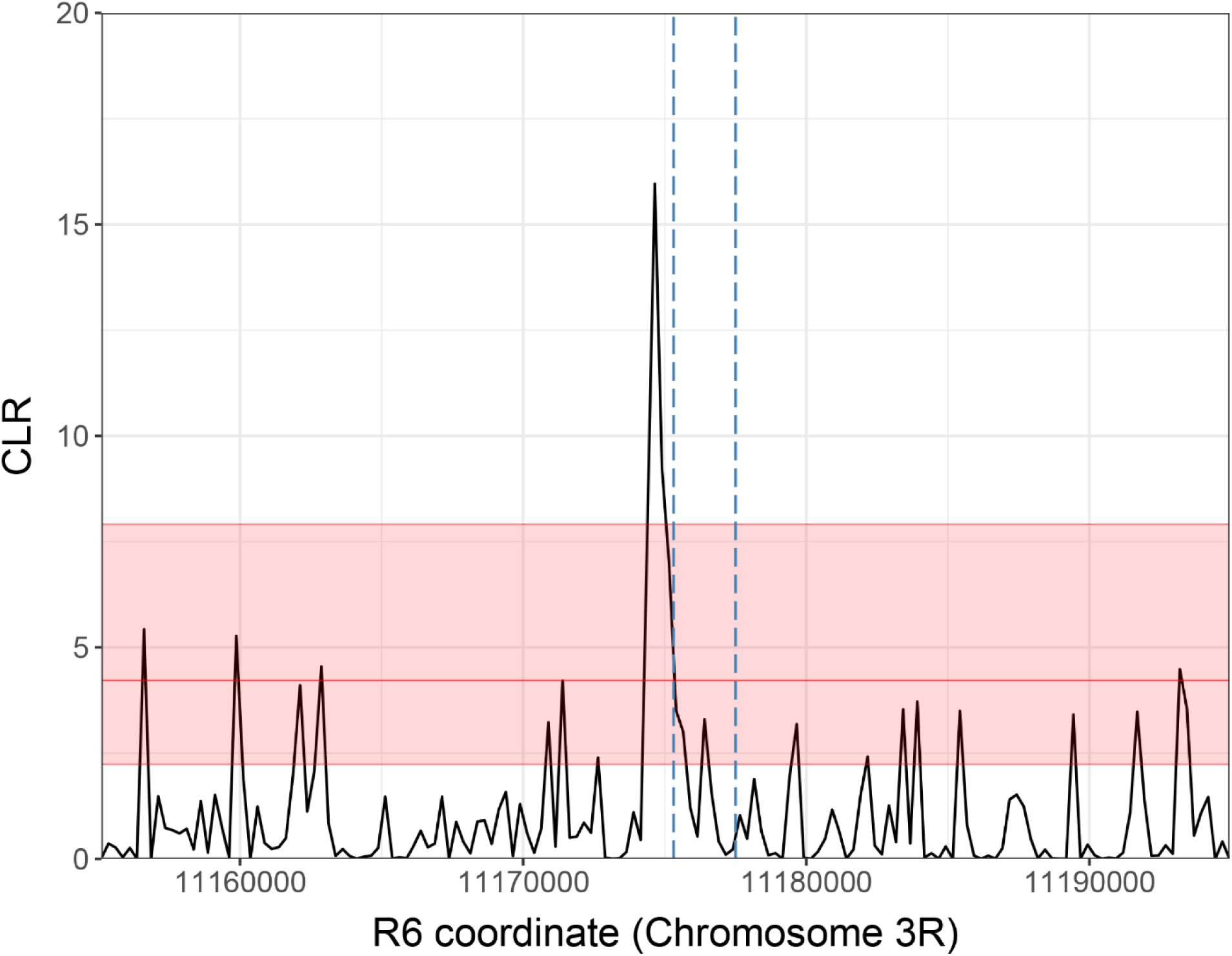
Distribution of CLR statistic (Nielsen, et al. 2005) at the genomic region containing the *Ugt86Dh* gene duplication in the sub-Saharan African populations, using SNPs called from (Lack, et al. 2015). The CLR peak falls immediately adjacent to the duplication. Given problems genotyping SNPs in duplicates, we do not actually expect the paralogous regions to be easily interpretable in a CLR framework. The shaded region represents the empirical 95% confidence interval for the maximum CLR values based on 100 neutral simulations using the SNPs present in this genomic region. Vertical blue lines indicate the breakpoints of the duplication discovered.

**Figure S19.**
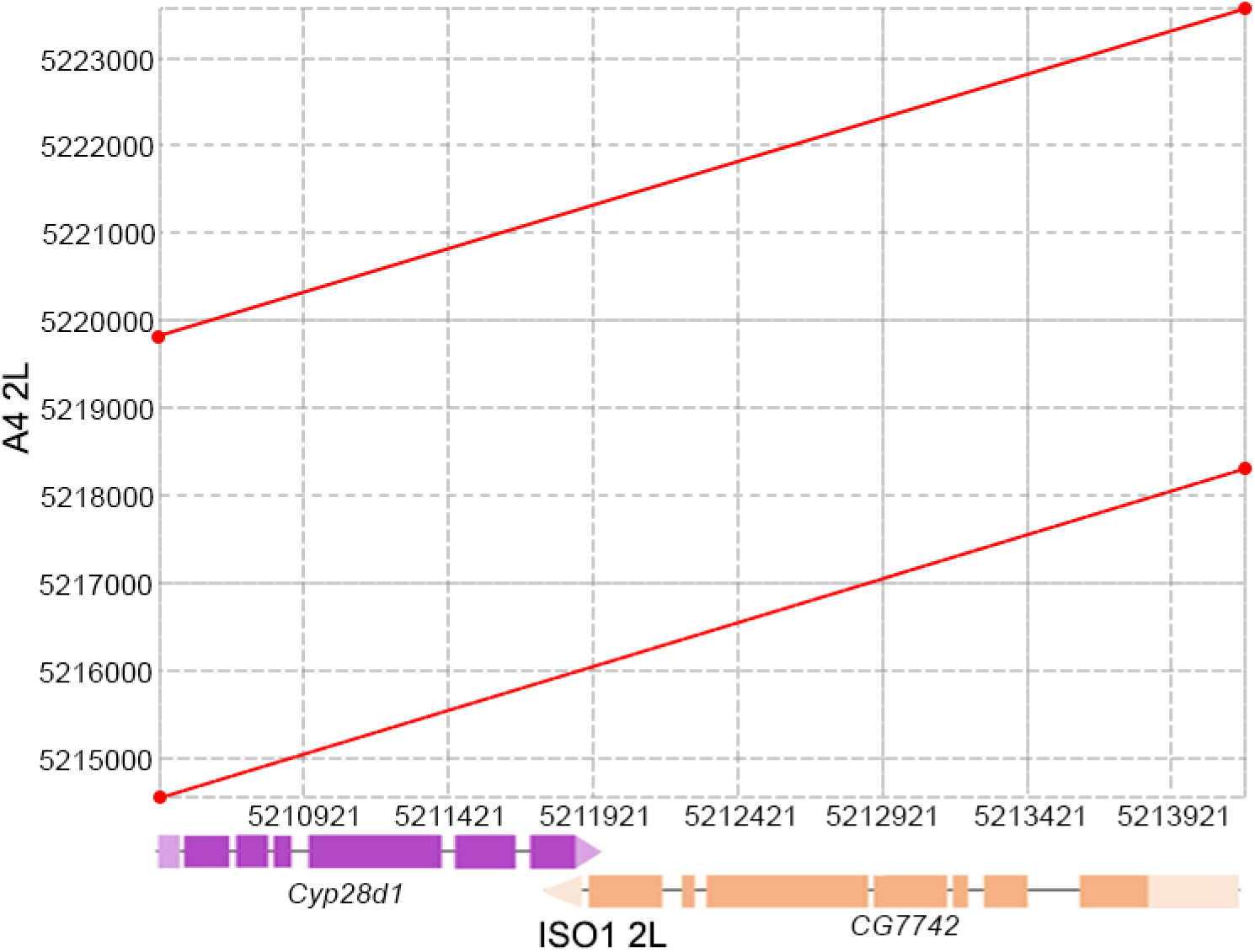
Dot plot between the ISO1 2L segment (5210421-5214176) containing *Cyp28d1* and *CG7742* and the region 5215000-5223000 on chromosome arm 2L in A4 (Y axis). The duplicates are shown as the two parallel lines spanning the entire ISO1 segment. The vertical space between the end of the bottom red line and the beginning of the top red line is due to the 1.5Kb fragment derived from an *Accord* element.

**>Supplementary Figure S20.**
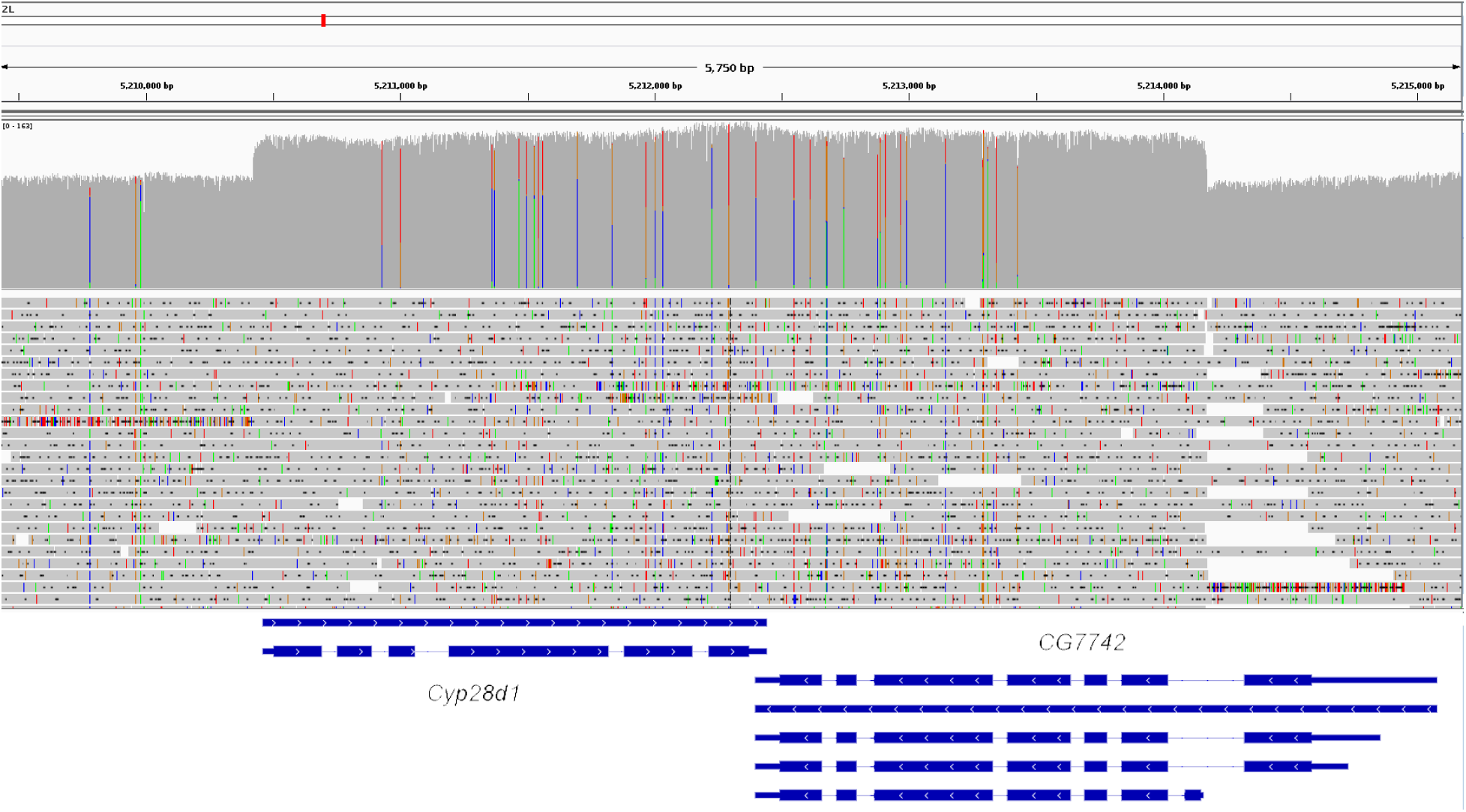
Alignment of A4 long reads to the ISO1genomic region (release 6) containing *Cyp28d1* (left) and *CG7742* (right). The region with higher read coverage (the grey hump) consists of the entire *Cyp28d1* and the smallest CG7742 isoform. The breakpoint on the right of the hump is due to the spacer sequence between the two copies.

**Figure S21.**
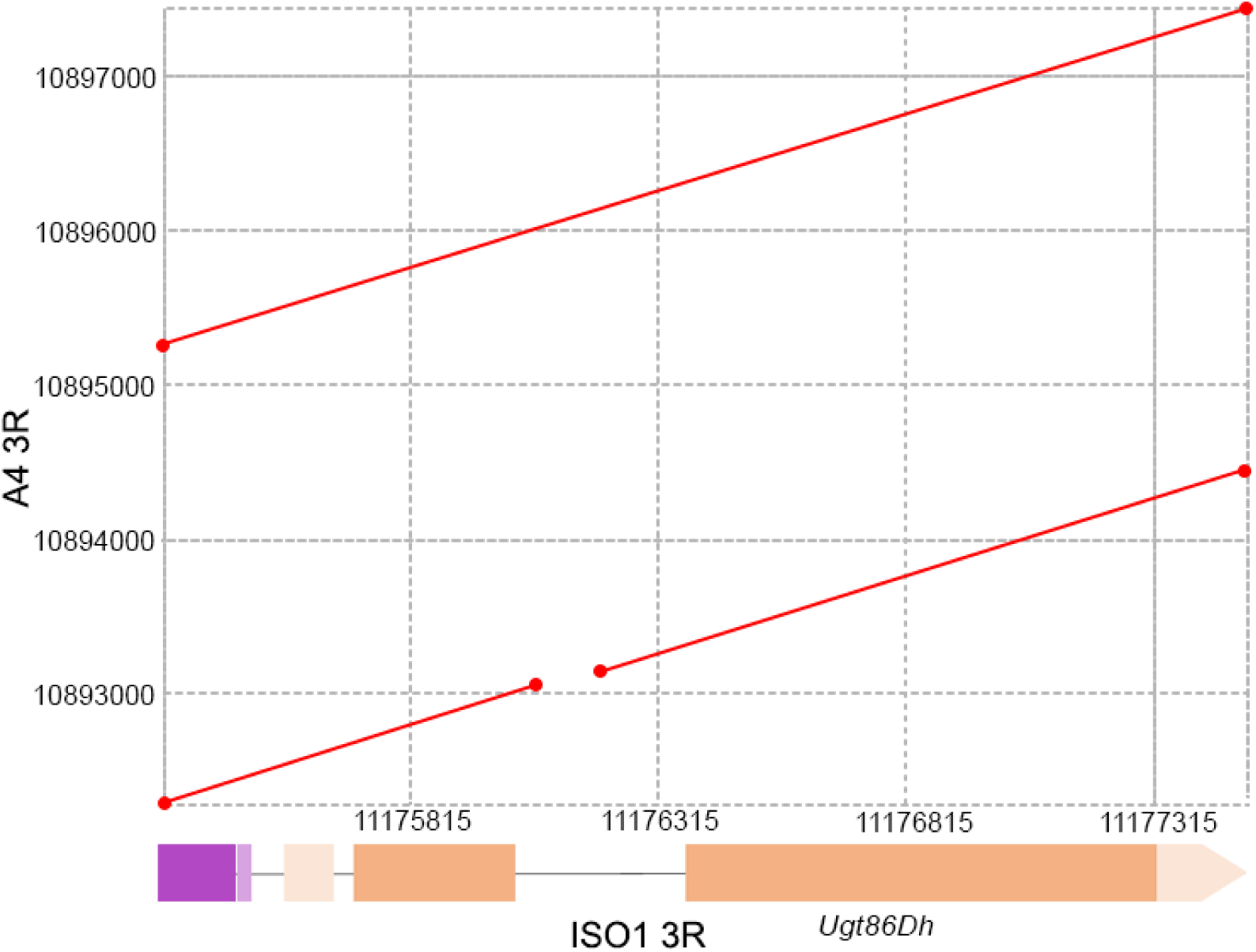
Dot plot between ISO1 genomic region 3R:11175315-11177501 and A4 genomic region 3R: 10892303-10897452 showing duplication of *Ugt86Dh* in A4. One of the copies in A4 has a deletion within the second intron.

**Figure S22.**
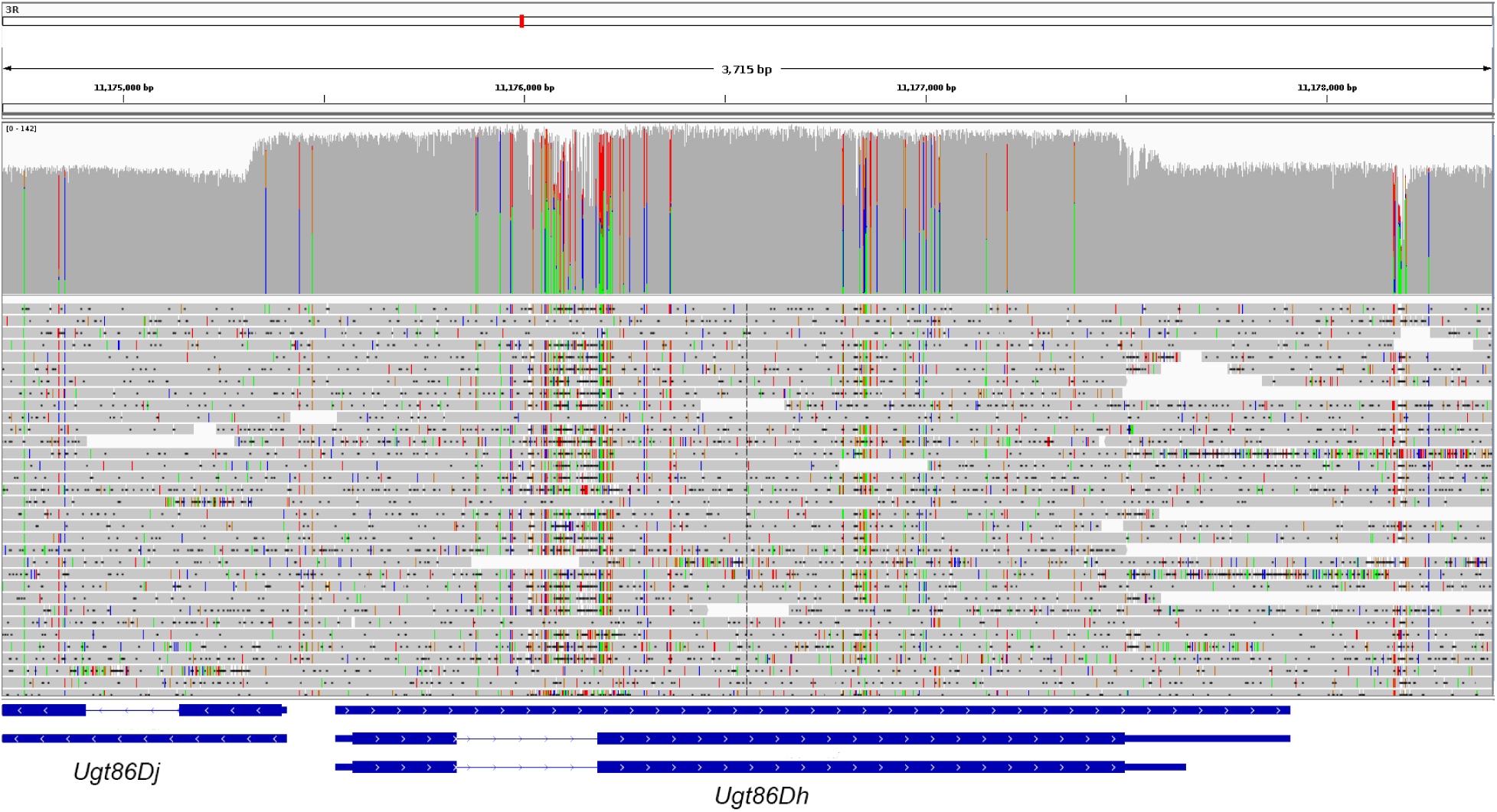
Alignment of A4 long reads to the ISO1genomic region containing a *Ugt86Dh* and a the first exon of *Ugt86Dj.* The blue rectangles at the bottom are the transcript annotations based on ISO1 release 6 coordinates. The region with greater read coverage (the grey hump) consists of the shorter isoform of *Ugt86Dh* and part of the *Ugt86Dj* first exon. As evidenced by the coverage drops along with gapped read alignment inside the hump, one of the A4 *Ugt86Dh* copies (*Ugt86Dh-d*) has a deletion inside an intron.

**Figure S23.**
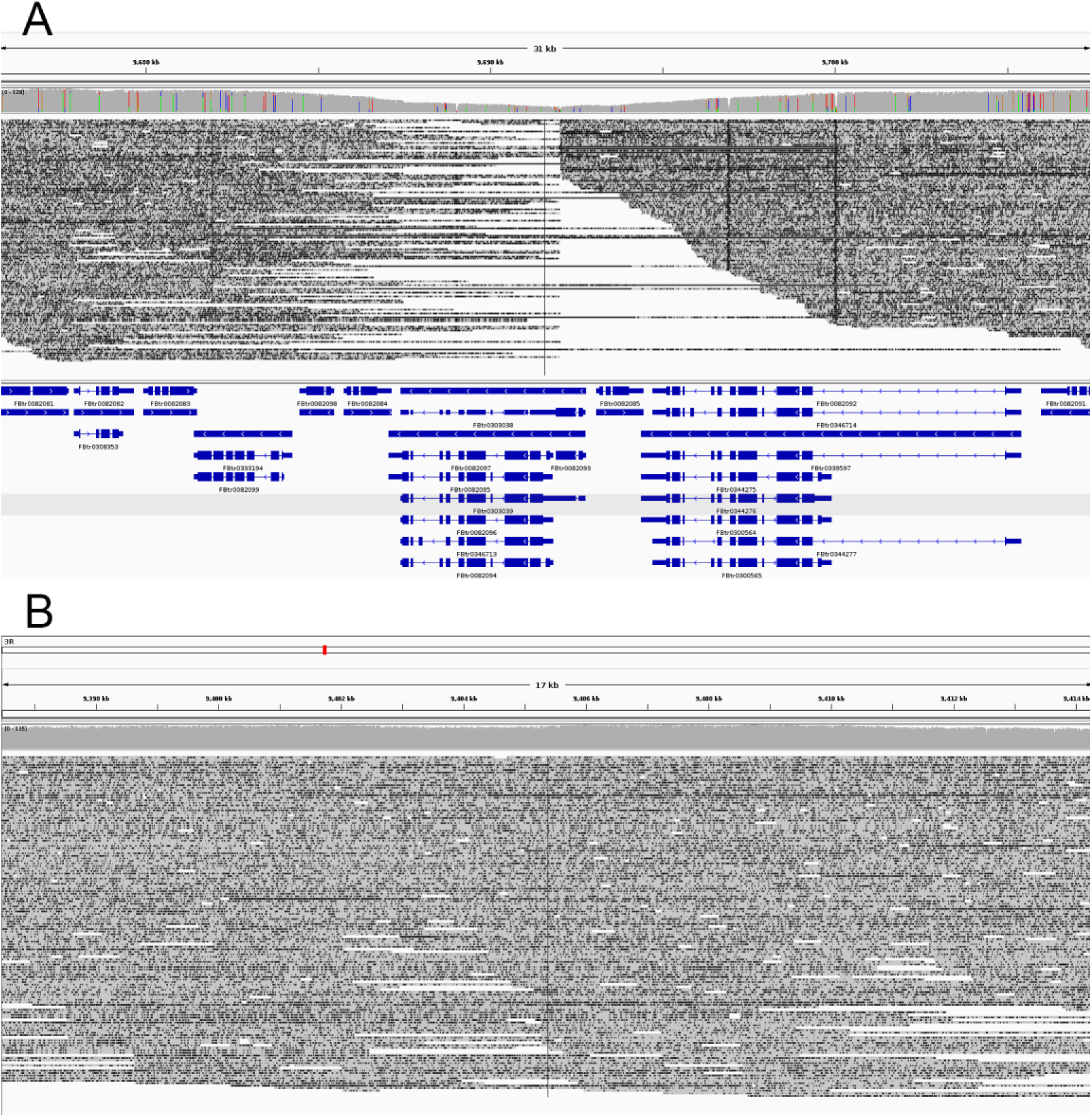
Absence of *p24-2* in A4. A) A4 long read coverage drops to zero at the genomic location of *p24-2* (FBtr0082093) in the ISO1 assembly, showing that A4 does not have *p24-2,* a gene claimed in(Chen, et al. 2010) as essential for *D. melanogaster.* B) A4 long reads aligned to the genomic region containing *Éclair* (*p24-2* paralog) in the A4 assembly, showing no abnormal read coverage.

**Figure S24.**
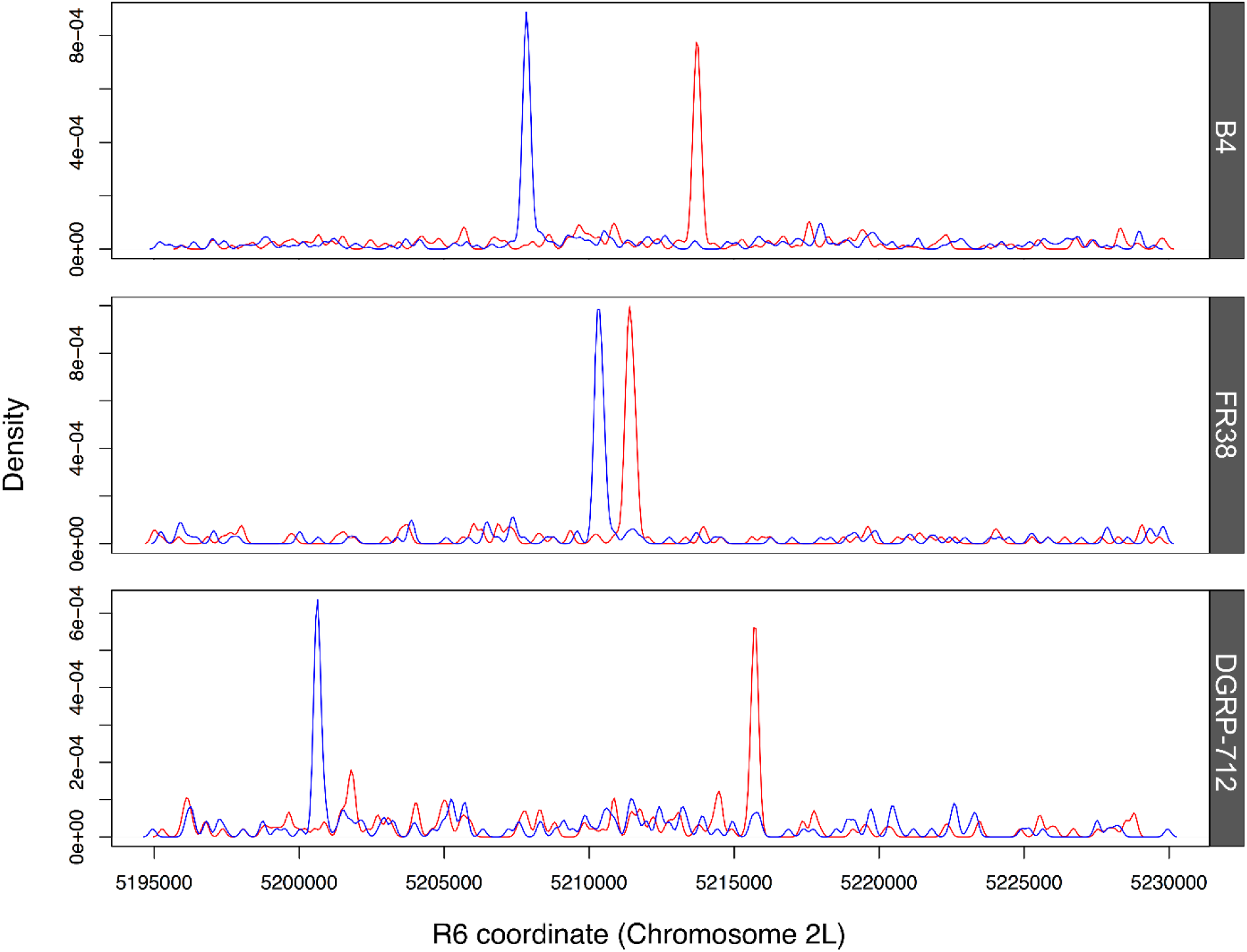
Divergent read pair coverage for several alleles at the *Cyp28d1* locus. Samples (top to bottom) are from Zimbabwe (DSPR), Riverside (DSPR), France (Bergman and Haddrill 2015), and North Carolina (Mackay, et al. 2012). Blue indicates reads aligning to the forward strand and red represents the reads aligning to the reverse strand.

**Table S1.** Comparison of assembly metrics between the A4 assembly and the release 6 of the reference assembly. The mitochondrial genome was excluded from both genomes and the Y-chromosome sequences were excluded from the ISO1 genome (the A4 genome has no Y-chromosome sequence, as it was assembled from reads derived from females).

| **Metric** | **ISO1** | **A4** |
| --- | --- | --- |
| **Assembly size (bp)** | 139,543,958 | 144,107,024 |
| **Contig N50 (bp)** | 21,485,538 | 22,302,559 |
| **Scaffold N50 (bp)** | 25,286,936 | 25,479,258 |
| **Number contigs** | 2,177 | 193 |
| **Number scaffolds** | 1,856 | 159 |
| **Complete BUSCO** | 2,625 | 2,621 |
| Single Copy | 2,492 | 2,495 |
| Duplicated | 133 | 126 |
| **Fragmented BUSCO** | 29 | 33 |
| **Missing BUSCO** | 21 | 21 |

**Table S2.** Chromosomal segments used for analyzing functional significance of the structural mutations (heterochromatic sequences are excluded following the coordinates in(Hoskins, et al. 2002)

| **Chromosome** | **Start** | **End** |
| --- | --- | --- |
| 2L | 1 | 22200000 |
| 2R | 4700000 | 25479258 |
| 3L | 1 | 23400000 |
| 3R | 3000000 | 31815305 |
| X | 1 | 21900000 |

Table S3. Coordinates of the CNVs, indels, and inversions in A4 and ISO1. (Table S2.xlsx)

Table S4. CNVs and indels for chromosome arms 2L as called by various CNV (*Pindel*(*Ye,* et al. 2009), *Pecnv*(*Rogers,* et al. 2014), and TE insertion(Cridland and Thornton 2010) calling softwares. (Table S4.xlsx)

Table S5. Summary of putative TE introns. (Table S5.xlsx)

Table S6. Genes mutated by non-TE indels in A4 (Table S6.xlsx).

Table S7. Expression changes in genes in A4 with increased copy number. (Table S7.xlsx)

**Table S8.**
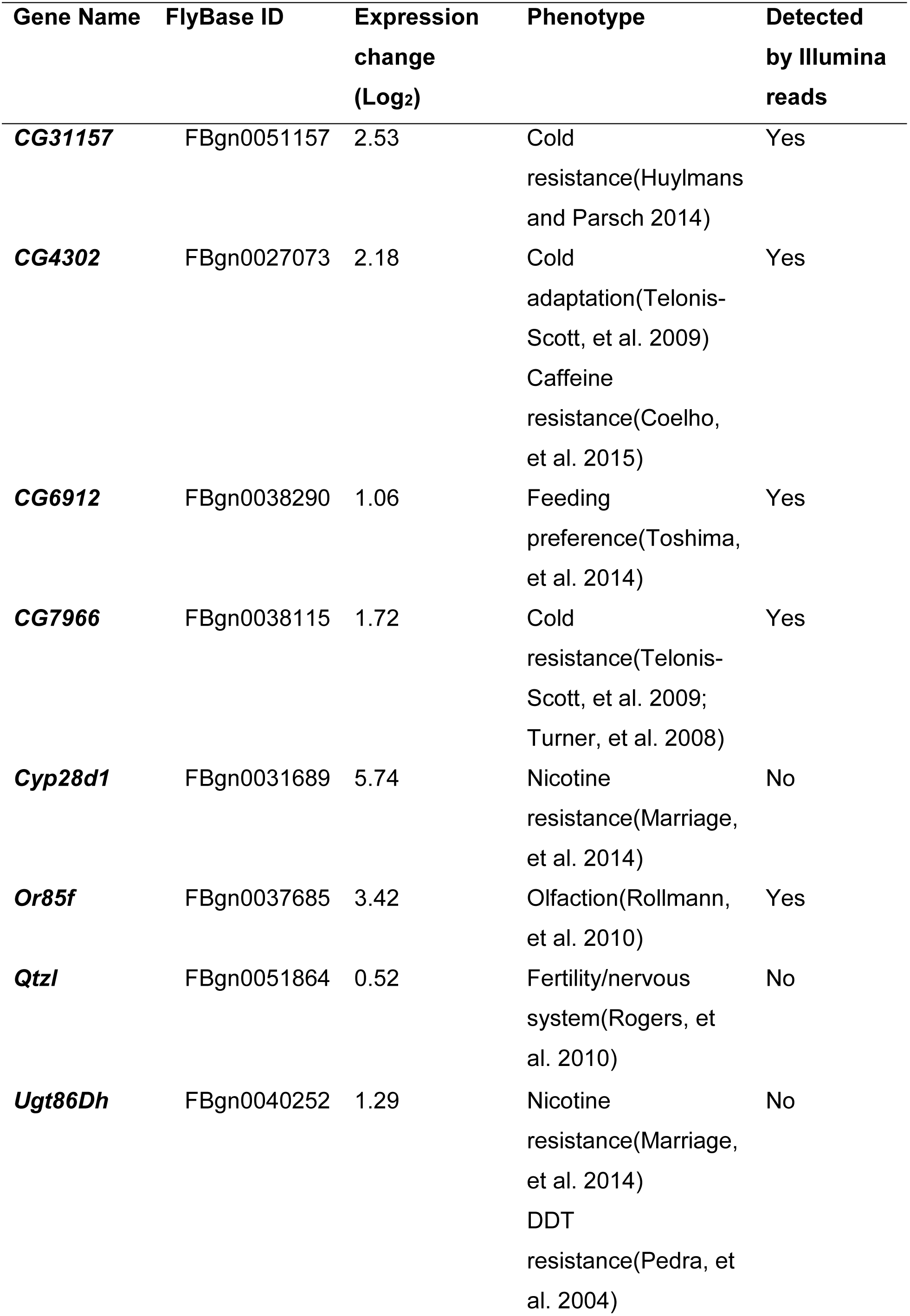
A4 duplicated genes with putative adaptive role.

## Acknowledgement

We thank Luna Thanh Ngo, Joshua Yan, and Allan Yue for help with fly maintenance and SV analysis. We also thank Jaaved Mohammed for kindly providing the multiple sequence alignment of the Drosophila species group. Finally, we would also like to thank Anthony Long, Brandon Gaut, Kevin Thornton for comments on the manuscript.

## References

2010. The human genome at ten. Nature 464: 649–650. doi: 10.1038/464649a

Abyzov A, Urban AE, Snyder M, Gerstein M 2011. CNVnator: An approach to discover, genotype, and characterize typical and atypical CNVs from family and population genome sequencing. Genome Research 21: 974–984. doi: 10.1101/gr.114876.110

Alkan C, Coe BP, Eichler EE 2011a. Genome structural variation discovery and genotyping. Nat Rev Genet 12: 363–376. doi: 10.1038/nrg2958

Alkan C, Sajjadian S, Eichler EE 2011b. Limitations of next-generation genome sequence assembly. Nature Methods 8: 61–65.

Bergman CM, Haddrill PR 2015. Strain-specific and pooled genome sequences for populations of Drosophila melanogaster from three continents. F1000Res 4: 31. doi: 10.12688/f1000research.6090.1

Berlin K, Koren S, Chin CS, Drake JP, Landolin JM, Phillippy AM 2015. Assembling large genomes with single-molecule sequencing and locality-sensitive hashing. Nat Biotechnol 33: 623–630. doi: 10.1038/nbt.3238

Chaisson MJ, Tesler G 2012. Mapping single molecule sequencing reads using basic local alignment with successive refinement (BLASR): application and theory. Bmc Bioinformatics 13. doi: Artn 238 10.1186/1471-2105-13-238

Chakraborty M, Baldwin-Brown JG, Long AD, Emerson JJ 2016. Contiguous and accurate de novo assembly of metazoan genomes with modest long read coverage. Nucleic Acids Res. doi: 10.1093/nar/gkw654

Chen S, Zhang YE, Long M 2010. New genes in Drosophila quickly become essential. Science 330: 1682–1685. doi: 10.1126/science.1196380

Chung H, Bogwitz MR, McCart C, Andrianopoulos A, Ffrench-Constant RH, Batterham P, Daborn PJ 2007. Cis-regulatory elements in the Accord retrotransposon result in tissue-specific expression of the Drosophila melanogaster insecticide resistance gene Cyp6g1. Genetics 175: 1071–1077. doi: 10.1534/genetics.106.066597

Coelho A, Fraichard S, Le Goff G, Faure P, Artur Y, Ferveur JF, Heydel JM 2015. Cytochrome P450- Dependent Metabolism of Caffeine in Drosophila melanogaster. Plos One 10. doi: UNSP e011732810.1371/journal.pone.0117328

Cridland JM, Macdonald SJ, Long AD, Thornton KR 2013. Abundance and distribution of transposable elements in two Drosophila QTL mapping resources. Mol Biol Evol 30: 2311–2327. doi: 10.1093/molbev/mst129

Cridland JM, Thornton KR 2010. Validation of rearrangement break points identified by paired-end sequencing in natural populations of Drosophila melanogaster. Genome Biol Evol 2: 83–101. doi: 10.1093/gbe/evq001

Cridland JM, Thornton KR, Long AD 2015. Gene expression variation in Drosophila melanogaster due to rare transposable element insertion alleles of large effect. Genetics 199: 85–93. doi: 10.1534/genetics.114.170837

DeGiorgio M, Huber CD, Hubisz MJ, Hellmann I, Nielsen R 2016. SweepFinder2: increased sensitivity, robustness and flexibility. Bioinformatics 32: 1895–1897. doi: 10.1093/bioinformatics/btw051

dos Santos G, Schroeder AJ, Goodman JL, Strelets VB, Crosby MA, Thurmond J, Emmert DB, Gelbart WM, FlyBase C 2015. FlyBase: introduction of the Drosophila melanogaster Release 6 reference genome assembly and large-scale migration of genome annotations. Nucleic Acids Res 43: D690–697. doi: 10.1093/nar/gku1099

Eichler EE, Flint J, Gibson G, Kong A, Leal SM, Moore JH, Nadeau JH 2010. Missing heritability and strategies for finding the underlying causes of complex disease. Nature Reviews Genetics 11: 446–450.

Emerson JJ, Cardoso-Moreira M, Borevitz JO, Long M 2008. Natural selection shapes genome-wide patterns of copy-number polymorphism in Drosophila melanogaster. Science 320: 1629–1631. doi: 10.1126/science.1158078

Frazer KA, Murray SS, Schork NJ, Topol EJ 2009. Human genetic variation and its contribution to complex traits. Nat Rev Genet 10: 241–251. doi: 10.1038/nrg2554

Gamazon ER, Nicolae DL, Cox NJ 2011. A Study of CNVs As Trait-Associated Polymorphisms and As Expression Quantitative Trait Loci. Plos Genetics 7. doi: ARTN e1001292 10.1371/journal.pgen.1001292

Glendinning JI 2002. How do herbivorous insects cope with noxious secondary plant compounds in their diet? Entomologia Experimentalis Et Applicata 104: 15–25. doi: DOI 10.1046/j.1570-7458.2002.00986.x

Henrichsen CN, Vinckenbosch N, Zollner S, Chaignat E, Pradervand S, Schutz F, Ruedi M, Kaessmann H, Reymond A 2009. Segmental copy number variation shapes tissue transcriptomes. Nature Genetics 41: 424–429.

Hoskins RA, Carlson JW, Wan KH, Park S, Mendez I, Galle SE, Booth BW, Pfeiffer BD, George RA, Svirskas R, Krzywinski M, Schein J, Accardo MC, Damia E, Messina G, Mendez-Lago M, de Pablos B, Demakova OV, Andreyeva EN, Boldyreva LV, Marra M, Carvalho AB, Dimitri P, Villasante A, Zhimulev IF, Rubin GM, Karpen GH, Celniker SE 2015. The Release 6 reference sequence of the Drosophila melanogaster genome. Genome Research 25: 445–458.

Hoskins RA, Smith CD, Carlson JW, Carvalho AB, Halpern A, Kaminker JS, Kennedy C, Mungall CJ, Sullivan BA, Sutton GG, Yasuhara JC, Wakimoto BT, Myers EW, Celniker SE, Rubin GM, Karpen GH 2002. Heterochromatic sequences in a Drosophila whole-genome shotgun assembly. Genome Biol 3: RESEARCH0085.

Huber CD, DeGiorgio M, Hellmann I, Nielsen R 2016. Detecting recent selective sweeps while controlling for mutation rate and background selection. Molecular Ecology 25: 142–156.

Huddleston J, Eichler EE 2016. An Incomplete Understanding of Human Genetic Variation. Genetics 202: 1251–1254.

Hudson RR 2002. Generating samples under a Wright-Fisher neutral model of genetic variation. Bioinformatics 18: 337–338.

Huylmans AK, Parsch J 2014. Population- and sex-biased gene expression in the excretion organs of Drosophila melanogaster. G3 (Bethesda) 4: 2307–2315. doi: 10.1534/g3.114.013417

Kang J, Kim J, Choi KW 2011. Novel Cytochrome P450, cyp6a17, Is Required for Temperature Preference Behavior in Drosophila. Plos One 6. doi: ARTN e29800 10.1371/journal.pone.0029800

Khost DE, Eickbush DG, Larracuente AM 2016. Single molecule long read sequencing resolves the detailed structure of complex satellite DNA loci in <em>Drosophila melanogaster</em>. bioRxiv.

King EG, Kislukhin G, Walters KN, Long AD 2014. Using Drosophila melanogaster To Identify Chemotherapy Toxicity Genes. Genetics 198: 31–+. doi: 10.1534/genetics.114.161968

King EG, Merkes CM, McNeil CL, Hoofer SR, Sen S, Broman KW, Long AD, Macdonald SJ 2012. Genetic dissection of a model complex trait using the Drosophila Synthetic Population Resource. Genome Research 22: 1558–1566.

Kurtz S, Phillippy A, Delcher AL, Smoot M, Shumway M, Antonescu C, Salzberg SL 2004. Versatile and open software for comparing large genomes. Genome Biol 5: R12. doi: 10.1186/gb-2004-5-2-r12

Lack JB, Cardeno CM, Crepeau MW, Taylor W, Corbett-Detig RB, Stevens KA, Langley CH, Pool JE 2015. The Drosophila genome nexus: a population genomic resource of 623 Drosophila melanogaster genomes, including 197 from a single ancestral range population. Genetics 199: 1229–1241. doi: 10.1534/genetics.115.174664

Lam KK, LaButti K, Khalak A, Tse D 2015. FinisherSC: a repeat-aware tool for upgrading de novo assembly using long reads. Bioinformatics 31: 3207–3209. doi: 10.1093/bioinformatics/btv280

Langmead B, Salzberg SL 2012. Fast gapped-read alignment with Bowtie 2. Nature Methods 9: 357–U354. doi: 10.1038/nmeth.1923

Li H, Durbin R 2009. Fast and accurate short read alignment with Burrows-Wheeler transform. Bioinformatics 25: 1754–1760. doi: 10.1093/bioinformatics/btp324

Li H, Handsaker B, Wysoker A, Fennell T, Ruan J, Homer N, Marth G, Abecasis G, Durbin R, Genome Project Data P 2009. The Sequence Alignment/Map format and SAMtools. Bioinformatics 25: 2078–2079. doi: 10.1093/bioinformatics/btp352

Luque T, O'Reilly DR 2002. Functional and phylogenetic analyses of a putative Drosophila melanogaster UDP-glycosyltransferase gene. Insect Biochemistry and Molecular Biology 32: 1597–1604.

Mackay TFC, Richards S, Stone EA, Barbadilla A, Ayroles JF, Zhu DH, Casillas S, Han Y, Magwire MM, Cridland JM, Richardson MF, Anholt RRH, Barron M, Bess C, Blankenburg KP, Carbone MA, Castellano D, Chaboub L, Duncan L, Harris Z, Javaid M, Jayaseelan JC, Jhangiani SN, Jordan KW, Lara F, Lawrence F, Lee SL, Librado P, Linheiro RS, Lyman RF, Mackey AJ, Munidasa M, Muzny DM, Nazareth L, Newsham I, Perales L, Pu LL, Qu C, Ramia M, Reid JG, Rollmann SM, Rozas J, Saada N, Turlapati L, Worley KC, Wu YQ, Yamamoto A, Zhu YM, Bergman CM, Thornton KR, Mittelman D, Gibbs RA 2012. The Drosophila melanogaster Genetic Reference Panel. Nature 482: 173–178. doi: 10.1038/nature10811

MacMillan HA, Knee JM, Dennis AB, Udaka H, Marshall KE, Merritt TJS, Sinclair BJ 2016. Cold acclimation wholly reorganizes the Drosophila melanogaster transcriptome and metabolome. Sci Rep 6. doi: Artn 28999 10.1038/Srep28999

Manolio TA, Collins FS, Cox NJ, Goldstein DB, Hindorff LA, Hunter DJ, McCarthy MI, Ramos EM, Cardon LR, Chakravarti A, Cho JH, Guttmacher AE, Kong A, Kruglyak L, Mardis E, Rotimi CN, Slatkin M, Valle D, Whittemore AS, Boehnke M, Clark AG, Eichler EE, Gibson G, Haines JL, Mackay TFC, McCarroll SA, Visscher PM 2009. Finding the missing heritability of complex diseases. Nature 461: 747–753.

Marriage TN, King EG, Long AD, Macdonald SJ 2014. Fine-mapping nicotine resistance loci in Drosophila using a multiparent advanced generation inter-cross population. Genetics 198: 45–57.

McCarthy MI, Abecasis GR, Cardon LR, Goldstein DB, Little J, Ioannidis JP, Hirschhorn JN 2008. Genomewide association studies for complex traits: consensus, uncertainty and challenges. Nat Rev Genet 9: 356–369. doi: 10.1038/nrg2344

mod EC, Roy S, Ernst J, Kharchenko PV, Kheradpour P, Negre N, Eaton ML, Landolin JM, Bristow CA, Ma L, Lin MF, Washietl S, Arshinoff BI, Ay F, Meyer PE, Robine N, Washington NL, Di Stefano L, Berezikov E, Brown CD, Candeias R, Carlson JW, Carr A, Jungreis I, Marbach D, Sealfon R, Tolstorukov MY, Will S, Alekseyenko AA, Artieri C, Booth BW, Brooks AN, Dai Q, Davis CA, Duff MO, Feng X, Gorchakov AA, Gu T, Henikoff JG, Kapranov P, Li R, MacAlpine HK, Malone J, Minoda A, Nordman J, Okamura K, Perry M, Powell SK, Riddle NC, Sakai A, Samsonova A, Sandler JE, Schwartz YB, Sher N, Spokony R, Sturgill D, van Baren M, Wan KH, Yang L, Yu C, Feingold E, Good P, Guyer M, Lowdon R, Ahmad K, Andrews J, Berger B, Brenner SE, Brent MR, Cherbas L, Elgin SC, Gingeras TR, Grossman R, Hoskins RA, Kaufman TC, Kent W, Kuroda MI, Orr-Weaver T, Perrimon N, Pirrotta V, Posakony JW, Ren B, Russell S, Cherbas P, Graveley BR, Lewis S, Micklem G, Oliver B, Park PJ, Celniker SE, Henikoff S, Karpen GH, Lai EC, MacAlpine DM, Stein LD, White KP, Kellis M 2010. Identification of functional elements and regulatory circuits by Drosophila modENCODE. Science 330: 1787–1797. doi: 10.1126/science.1198374

Myers EW, Sutton GG, Delcher AL, Dew IM, Fasulo DP, Flanigan MJ, Kravitz SA, Mobarry CM, Reinert KH, Remington KA, Anson EL, Bolanos RA, Chou HH, Jordan CM, Halpern AL, Lonardi S, Beasley EM, Brandon RC, Chen L, Dunn PJ, Lai Z, Liang Y, Nusskern DR, Zhan M, Zhang Q, Zheng X, Rubin GM, Adams MD, Venter JC 2000. A whole-genome assembly of Drosophila. Science 287: 2196–2204.

Nielsen R, Williamson S, Kim Y, Hubisz MJ, Clark AG, Bustamante C 2005. Genomic scans for selective sweeps using SNP data. Genome Res 15: 1566–1575. doi: 10.1101/gr.4252305

Pedra JHF, McIntyre LM, Scharf ME, Pittendrigh BR 2004. Genom-wide transcription profile of field- and laboratory-selected dichlorodiphenyltrichloroethane (DDT)-resistant Drosophila. Proceedings of the National Academy of Sciences of the United States of America 101: 7034–7039. doi: 10.1073/pnas.0400580101

Petrov DA, Fiston-Lavier A-S, Lipatov M, Lenkov K, Gonzalez J 2011. Population genomics of transposable elements in Drosophila melanogaster. Molecular Biology and Evolution 28: 1633–1644.

Quinlan AR 2014. BEDTools: The Swiss-Army Tool for Genome Feature Analysis. Current protocols in bioinformatics / editoral board, Andreas D Baxevanis [et al] 47: 11 12 11–34. doi: 10.1002/0471250953.bi1112s47

Rebollo R, Miceli-Royer K, Zhang Y, Farivar S, Gagnier L, Mager DL 2012. Epigenetic interplay between mouse endogenous retroviruses and host genes. Genome Biol 13: R89. doi: 10.1186/gb-2012-13-10-r89

Rockman MV 2012. THE QTN PROGRAM AND THE ALLELES THAT MATTER FOR EVOLUTION: ALL THAT'S GOLD DOES NOT GLITTER. Evolution 66. doi: 10.1111/j.1558-5646.2011.01486.x

Rogers RL, Bedford T, Lyons AM, Hartl DL 2010. Adaptive impact of the chimeric gene Quetzalcoatl in Drosophila melanogaster. Proceedings of the National Academy of Sciences of the United States of America 107: 10943–10948. doi: 10.1073/pnas.1006503107

Rogers RL, Cridland JM, Shao L, Hu TT, Andolfatto P, Thornton KR 2014. Landscape of standing variation for tandem duplications in Drosophila yakuba and Drosophila simulans. Mol Biol Evol 31: 1750–1766. doi: 10.1093/molbev/msu124

Rollmann SM, Wang P, Date P, West SA, Mackay TFC, Anholt RRH 2010. Odorant Receptor Polymorphisms and Natural Variation in Olfactory Behavior in Drosophila melanogaster. Genetics 186: 687–U364. doi: 10.1534/genetics.110.119446

Schmidt JM, Good RT, Appleton B, Sherrard J, Raymant GC, Bogwitz MR, Martin J, Daborn PJ, Goddard ME, Batterham P, Robin C 2010. Copy number variation and transposable elements feature in recent, ongoing adaptation at the Cyp6g1 locus. PLoS Genet 6: e1000998. doi: 10.1371/journal.pgen.1000998

Simao FA, Waterhouse RM, Ioannidis P, Kriventseva EV, Zdobnov EM 2015. BUSCO: assessing genome assembly and annotation completeness with single-copy orthologs. Bioinformatics 31: 3210–3212. doi: 10.1093/bioinformatics/btv351

Stapleton M, Liao G, Brokstein P, Hong L, Carninci P, Shiraki T, Hayashizaki Y, Champe M, Pacleb J, Wan K, Yu C, Carlson J, George R, Celniker S, Rubin GM 2002. The Drosophila gene collection: identification of putative full-length cDNAs for 70% of D. melanogaster genes. Genome Res 12: 1294–1300. doi: 10.1101/gr.269102

Stranger BE, Forrest MS, Dunning M, Ingle CE, Beazley C, Thorne N, Redon R, Bird CP, de Grassi A, Lee C, Tyler-Smith C, Carter N, Scherer SW, Tavare S, Deloukas P, Hurles ME, Dermitzakis ET 2007. Relative impact of nucleotide and copy number variation on gene expression phenotypes. Science 315: 848–853. doi: 10.1126/science.1136678

Swinburne IA, Silver PA 2008. Intron delays and transcriptional timing during development. Dev Cell 14: 324–330. doi: 10.1016/j.devcel.2008.02.002

Telonis-Scott M, Hallas R, McKechnie SW, Wee CW, Hoffmann AA 2009. Selection for cold resistance alters gene transcript levels in Drosophila melanogaster. Journal of Insect Physiology 55: 549–555. doi: 10.1016/j.jinsphys.2009.01.010

Thorvaldsdottir H, Robinson JT, Mesirov JP 2013. Integrative Genomics Viewer (IGV): high-performance genomics data visualization and exploration. Brief Bioinform 14: 178–192. doi: 10.1093/bib/bbs017

Toshima N, Hara C, Scholz CJ, Tanimura T 2014. Genetic variation in food choice behaviour of amino acid-deprived Drosophila. Journal of Insect Physiology 69: 89–94. doi: 10.1016/j.jinsphys.2014.06.019

Trapnell C, Roberts A, Goff L, Pertea G, Kim D, Kelley DR, Pimentel H, Salzberg SL, Rinn JL, Pachter L 2012. Differential gene and transcript expression analysis of RNA-seq experiments with TopHat and Cufflinks. Nature Protocols 7: 562–578. doi: 10.1038/nprot.2012.016

Turner TL, Levine MT, Eckert ML, Begun DJ 2008. Genomic analysis of adaptive differentiation in Drosophila melanogaster. Genetics 179: 455–473. doi: 10.1534/genetics.107.083659

Walker BJ, Abeel T, Shea T, Priest M, Abouelliel A, Sakthikumar S, Cuomo CA, Zeng Q, Wortman J, Young SK, Earl AM 2014. Pilon: an integrated tool for comprehensive microbial variant detection and genome assembly improvement. Plos One 9: e112963. doi: 10.1371/journal.pone.0112963

Wray NR, Yang J, Hayes BJ, Price AL, Goddard ME, Visscher PM 2013. Pitfalls of predicting complex traits from SNPs. Nat Rev Genet 14: 507–515. doi: 10.1038/nrg3457

Ye C, Hill CM, Wu S, Ruan J, Ma ZS 2016. DBG2OLC: Efficient Assembly of Large Genomes Using Long Erroneous Reads of the Third Generation Sequencing Technologies. Sci Rep 6: 31900. doi: 10.1038/srep31900

Ye K, Schulz MH, Long Q, Apweiler R, Ning Z 2009. Pindel: a pattern growth approach to detect break points of large deletions and medium sized insertions from paired-end short reads. Bioinformatics 25: 2865–2871. doi: 10.1093/bioinformatics/btp394

